# A peptide-centric local stability assay to unveil protein targets of diverse ligands

**DOI:** 10.1101/2023.10.17.562693

**Authors:** Kejia Li, Shijie Chen, Keyun Wang, Yan Wang, Zheng Fang, Jiawen Lyu, Haiyang Zhu, Yanan Li, Ting Yu, Feng Yang, Xiaolei Zhang, Siqi Guo, Chengfei Ruan, Jiahua Zhou, Qi Wang, Cheng Luo, Mingliang Ye

## Abstract

While tremendous progress has been made in chemical proteomics for identifying protein-ligand interactions, it remains challenging for proteome-wide identification of ligand-binding regions without modifying the ligands. Here, we discovered that “disruptive trypsinization” amplifies the readout of ligand-induced protein local stability shifts, and explored this notion in developing “peptide-centric local stability assay” (PELSA), a modification-free approach which achieves unprecedented sensitivity in proteome-wide target identification and binding-region determination. We demonstrate the versatility of PELSA by investigating the interactions across various biological contexts including drug-target interactions, metabolism, epitope mapping, metal proteomics, and post-translational modification recognition. A PELSA study of the oncometabolite R2HG revealed functional insights about its targets and pathogenic processes in both cancer and immune cells. Thus, beyond offering users unprecedented sensitivity for characterizing diverse target-ligand interactions, PELSA supports informative screening and hypothesis generation studies throughout life science.

## INTRODUCTION

The biochemical functions of proteins invariably involve interactions with ligands of some type, which act as enzyme substrates or inhibitors, signaling molecules, allosteric modulators, structural anchors, etc. Monitoring protein-ligand interactions is thus essential for comprehending various aspects of life science, including drug mechanisms of action, regulatory processes in cellular metabolism and signaling, and the functions of uncharacterized proteins^1, 2^. Additionally, knowledge of the ligand-binding regions holds immense value for structure-based drug design^3^ and biological hypothesis generation^4^.

Modification-based methods which rely on chemical modifications of the ligands to capture ligand-binding proteins and binding regions^5, 6^ require extensive chemical synthesis and may be not applicable for ligands that lack suitable sites for chemical modification^7^. Previously reported modification-free methods^8^, including CEllular Thermal Shift Assay (CETSA)^9^ and Thermal Proteome Profiling (TPP)^10^ bypass the need of ligand modification but do not support the identification of specific ligand-binding regions in target proteins. Paola et al. developed LiP-MS (limited proteolysis coupled with mass spectrometry)^11^, which can identify ligand-binding proteins and binding regions in the cell lysates of microbial organisms. Despite the advancements brought by LiP-MS and the subsequently developed LiP-Quant^12^ (a dose-response version of LiP-MS tailored for complex human cell lysates), their capacity for target identification remains limited^12^.

Here, we propose disruptive trypsinization to directly generate MS-detectable peptides from native proteins to represent protein local stability. This digestion scheme in couple with a simple separation procedure largely reduces the complexity of peptide samples and, crucially, amplifies the readout of ligand-induced protein local stability shifts. Based on this observation, we established a method we term PEptide-centric Local Stability Assay (PELSA) that enables sensitive identification of target proteins while also preserving extensive binding-region information. We demonstrate that PELSA achieves unprecedented sensitivity in revealing ligand-binding proteins through extensive comparisons against alternative methods. For example, PELSA with one drug dose and one digesting condition identified 12-fold more kinase targets for a pan-kinase inhibitor than LiP-Quant using seven drug doses^12^, and 2.4-fold more targets than TPP using ten temperatures^10^. We further demonstrate the wide application scope and excellent performance of PELSA in studies of drug promiscuity, molecular glue, epitopes, recognition domains for post-translational modifications, metal proteomics, and metabolite sensing and signaling.

## RESULTS

### Disruptive trypsinization amplifies the readout of ligand-induced protein local stability shifts

Binding with a ligand can increase the stability of the ligand-binding region of a protein^13, 14^. The stability of a protein can be measured by its protease susceptibility^15,16^. Therefore, when native proteins are partially digested into small peptides, the abundance of every individual peptide should represent a measurement of the stability of the region in which it is located. We speculate these directly generated peptides could be used for investigating ligand-induced protein local stability shifts. Because these small peptides can be easily separated from the undigested large counterparts (e.g., through differences in molecular weight), the complexity of the resulted peptide mixture will be largely reduced (compared to LiP-MS^11, 12^, a two-step digestion scheme for ligand-binding protein identification), which could enable detection of a rich array of peptides that are informative regarding ligand binding.

Pursuing this, we used trypsinization with a high E/S ratio (enzyme/substrate, wt/wt) for a short time (*i.e*., 1 min) to partially digest native proteins into small peptides. We used trypsin because tryptic peptides are optimal for shotgun proteomics analysis^17^, and used a high E/S ratio to enable generation of a large number of small tryptic peptides. We term this digestion scheme as “disruptive trypsinization”, as trypsinization also functions here as a denaturant to destroy the protein structures to facilitate small peptide generation. The generated tryptic peptides are subsequently enriched by removing large, partially digested protein segments through a filter unit, followed by proteomics analysis.

To test if our procedure could identify more peptides that are informative regarding ligand binding than existing LiP-MS methods, we worked with HeLa cell lysates and two well-studied drugs: Methotrexate (MTX) targeting DHFR^18^ and SHP099 targeting PTPN11^19^. The comparison was performed between our procedure with an E/S ratio of 1:2 (trypsin: substrate, wt/wt) and the LiP-MS approach with an initial brief digestion at an E/S ratio of 1:100 (proteinase K: substrate, wt/wt) followed by complete trypsin digestion under denaturing conditions^12^. Note that the initial data quality assessment confirmed the proper operations of LiP-MS in our study (**Extended Data Fig. 1a,b**).

Gratifyingly, both the data from MTX and SHP099 experiments showed that our procedure identified many more peptides showing statistically significant abundance changes (Bayes t-test p < 0.01, |log_2_FC| > 0.3) on target proteins than LiP-MS (**Fig. 1a, Extended Data Fig. 1c**; 12 and 21 versus 6 and 4). We defined these peptides as “ligand-responsive target” (LRT) peptides. Strikingly, we observed that the LRT peptides displayed a remarkably larger “readout” (*i.e.*, magnitude of abundance fold changes) in our procedure than in LiP-MS (**Fig. 1b**; medians of |log_2_FC|: 4.07 and 3.35 versus 0.95 and 0.52). It bears emphasis that only peptides from the ligand-binding domains displayed an amplified readout when using disruptive trypsinization, whereas peptides from the unbound regions remained no abundance changes upon ligand treatment (**Fig. 1c**). The amplified readout may be because disruptive trypsinization is a continuous multi-stage proteolysis process, in which the abundance difference of the peptides generated from the ligand-binding regions between bound and unbound states, reflects an accumulation of differences in the rates of multi-stage proteolysis (**Extended Data Fig. 2**). Benefiting from the amplified readout, disruptive trypsinization yielded more accurate binding region data relative to LiP-MS (**Extended Data Fig. 3a-c**).

**Fig. 1.**
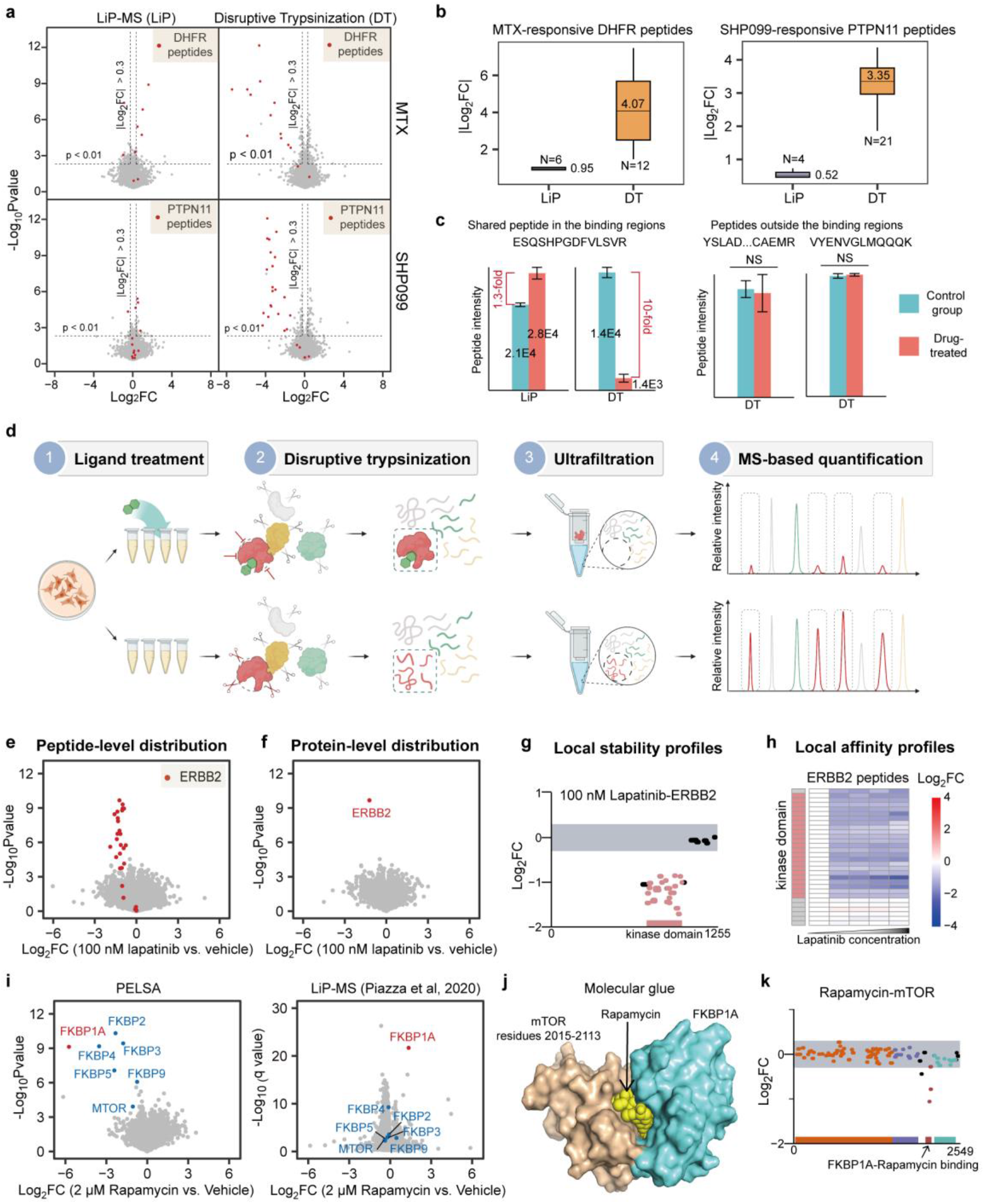
Establishment of PELSA. **a,** Volcano plot visualizations of all peptides generated by LiP-MS (LiP) or Disruptive Trypsinization (DT) of HeLa lysates exposed to 10 μM methotrexate (MTX) or 10 μM SHP099 (four lysate replicates per experiment). **b,** Comparing readouts of the ligand-responsive target (LRT) peptides generated by LiP-MS and disruptive trypsinization. Central line in the box shows the median (labeled), box boundaries indicate the upper and lower interquartile range (IQR), and whiskers correspond to most extreme values, or to 1.5-fold IQR if the extreme values are above this cutoff. **c,** Left: the peptide shared in two digestion schemes and from SHP099-binding domains, display amplified readout when using disruptive trypsinization. Right: two peptides located outside the SHP099-binding region remained unchanged by SHP099 treatment in disruptive trypsinization. Four replicates (mean ±S.D.). NS, not significant. **d**, Workflow of PELSA. **e,** Volcano plot visualization of all peptides from a PELSA analysis of BT474 lysates exposed to 100 nM lapatinib. **f**, Volcano plot as in (**e**) but on the protein-level. **g**, Local stability profiles to reveal ligand-binding regions. The upper and lower boundaries of the grey shaded area represent log_2_FCs of 0.3 and -0.3, respectively. **h,** Local affinity profiles to reveal the local binding affinity of a ligand. Heat map representation of log_2_ peptide fold changes of ERBB2 with increasing lapatinib concentrations (0 nM, 100 nM, 1 μM, 10 μM, and 100 μM). **i,** Volcano plot visualizations of all proteins from a PELSA analysis or a published LiP-MS analysis^12^ of HeLa lysates exposed to 2 μM rapamycin. **j**, Complex structure of mTOR, rapamycin, and FKBP1A (PDB: 1FAP). **k,** Local stability profiles of mTOR for 2 μM rapamycin treatment.

In conclusion, we demonstrate that our procedure can identify a rich array of peptides that are informative regarding ligand binding. Crucially, we found that disruptive trypsinization can amplify the readout of ligand-induced protein local stability shifts. Based on these observations, we propose a peptide-centric local stability assay, or PELSA, to probe ligand-binding proteins and binding regions.

### The PELSA approach

In the PELSA workflow (**Fig. 1d**), proteome samples extracted from cell lysates under native conditions are incubated with an analyte ligand (exemplified by lapatinib^20^, a marketed inhibitor of a membrane protein, ERBB2) or vehicle, respectively. The two sample groups are then subjected to trypsinization with a high E/S ratio (enzyme/substrate, wt/wt) (*e.g.,* 1:2) for a short time (*e.g.*, 1 min) followed by removing any large, partially digested protein fragments with an ultrafiltration unit (molecular weight cutoff 10 kDa). The collected peptides are then analyzed by liquid chromatography-tandem mass spectrometry (LC-MS/MS) in data-independent acquisition (DIA) mode. The quantified peptides are compared between two groups (Bayes t-test analysis) (**Fig. 1e**), and the peptide with the lowest p value among all quantified peptides of the same protein is selected to represent its corresponding protein for target protein identification (**Fig. 1f**). Notably, out of 5866 proteins, we identified the known lapatinib target protein ERBB2 as the top target candidate (unless otherwise stated, target prioritization is ranked by -log_10_Pvalue). Mapping the quantified peptides to protein sequences generates local stability profiles (**Fig. 1g**), which reveal the protein regions responsive to the ligand binding. Consistent with the previous knowledge that lapatinib binds ERBB2 via its kinase domain^20^, the PELSA local stability profile data showed that the ligand-responsive peptides detected for ERBB2 were all from the kinase domain (**Fig. 1g**). The dose-dependent local stability changes can also be assessed when PELSA experiments are performed using multiple ligand doses. Since the local stability changes of the target protein are dependent on the ligand occupancy, the dose that produces the half-maximal stability changes reflects the local binding affinity of the ligand for the corresponding protein segment. Hence, we termed the dose-response local stability changes as “local affinity profiles” (**Fig. 1h**).

We next applied PELSA to investigate the target proteins of rapamycin, an inhibitor of multiple FKBP family proteins^21^. A previous LiP-MS study successfully identified FKBP1A as a rapamycin-binding protein^12^. Initially, we applied PELSA to investigate rapamycin-binding proteins under experimental conditions identical to those reported in the LiP-MS paper (*i.e.*, HeLa cell lysates, 2 µM rapamycin). Consistent with the amplified readouts observed for DHFR and PTPN11, PELSA generated a 25-fold larger readout for FKBP1A than LiP-MS (53.4 versus 2.54) (**Fig. 1i**). Beyond FKBP1A, PELSA identified five additional FKBP family proteins as rapamycin-binding proteins (**Fig. 1i**), which failed to recognize as rapamycin-binding proteins in LiP-MS due to no detectable fold changes (**Fig. 1i**). We further demonstrated these additional FKBP family proteins has a low target occupancy under 2 µM rapamycin treatment **(Extended Data Fig. 3d).** These results suggested that the amplified readout equips PELSA with the sensitivity to identify low stoichiometry binding events in the cellular context.

PELSA can identify not only ligand-binding regions located on a single protein, but also those that span two proteins. Rapamycin can act as a molecular glue between FKBP1A and mTOR (**Fig. 1j**), and the FKBP1A-rapamycin complex binds to a small segment of mTOR (residues 2015-2113)^22^. Beyond successful determination of FKBP domains as rapamycin-binding regions on the identified FKBP family proteins (**Extended Data Fig. 3e)**, the PELSA local stability profiles accurately pinpointed the binding region of the FKBP1A-rapamycin complex on mTOR (**Fig. 1k**).

Besides DIA, other quantitative proteomics methods can also be used to quantify PELSA-generated peptides. For example, we coupled PELSA with the data-dependent acquisition (DDA) based cost-effective stable isotope dimethyl labeling^23^ to investigate the binding profiles of three HSP90 inhibitors with distinct structural similarities (**Extended Data Fig. 4a**). PELSA successfully identified HSP90 family proteins—and determined the known binding regions (*i.e.*, N-terminal ATP-binding domain)^24^ —for the three HSP90 inhibitors (**Extended Data Fig. 4b,c**). As expected, the structurally close inhibitors, geldanamycin and tanespimycin, shared more off-targets (**Extended Data Fig. 4b**). The unique off-targets identified for the structurally distinct inhibitor ganetespib, AKR1C2 and MAT2A, were also validated using thermal shift assay with purified proteins (**Extended Data Fig. 4d**), substantiating the reliability of PELSA for target identification. The dose-dependent PELSA analysis also yielded accurate binding affinity data (**Extended Data Fig. 4e**), aligning with results from microscale thermophoresis (MST) assay (**Extended Data Fig. 4f**).

Taken together, we demonstrate that PELSA enables efficient target identification, precise binding-region determination, and accurate binding affinity quantification on the proteome-wide scale.

### PELSA’s high sensitivity in target identification

Staurosporine, a pan-kinase inhibitor, has been investigated by LiP-Quant (the dose-dependent version of LiP-MS) in HeLa cell lysates^12^ and TPP in K562 cell lysates^10^. To compare the performance of PELSA for target identification against these popular modification-free methods, we screened the targets of staurosporine by PELSA in both lysates of HeLa and K562 cells and compared our results with the published datasets of LiP-Quant and TPP.

Using a true positive rate (TPR, defined as the percentage of kinase targets in candidate targets) cutoff of 80% (**Supplementary Discussion**), PELSA with one staurosporine dose yielded 120/143 (kinases/candidates) and 108/135 (kinases/candidates) in K562 and HeLa cell lysates, respectively (**Fig. 2a,b** **and Supplementary Table 1**). By contrast, a LiP-Quant analysis of staurosporine in HeLa cell lysates identified 20 kinase targets (TPR of 40%)^12^, and 9 kinase targets were identified when the identical criterion—TPR cutoff of 80%—was applied (**Fig. 2b** and **Supplementary Table 1**), albeit with 7 drug doses and a superior LC-MS/MS analysis depth reflected by more quantified peptides^12^ and higher protein sequence coverages (**Fig. 2c**). In line with our observations in MTX, SHP099, and rapamycin experiments, the overlapped kinase targets displayed much larger readouts in PELSA than in LiP-Quant (median values of |log_2_FC|: 2.2 versus 0.75) (**Fig. 2d**). Since in PELSA the peptides that are most relevant to ligand binding are enriched, PELSA requires lower protein sequence coverages than LiP-Quant for successful identification of target proteins (**Fig. 2c**), which also leads to the high sensitivity of PELSA.

**Fig. 2.**
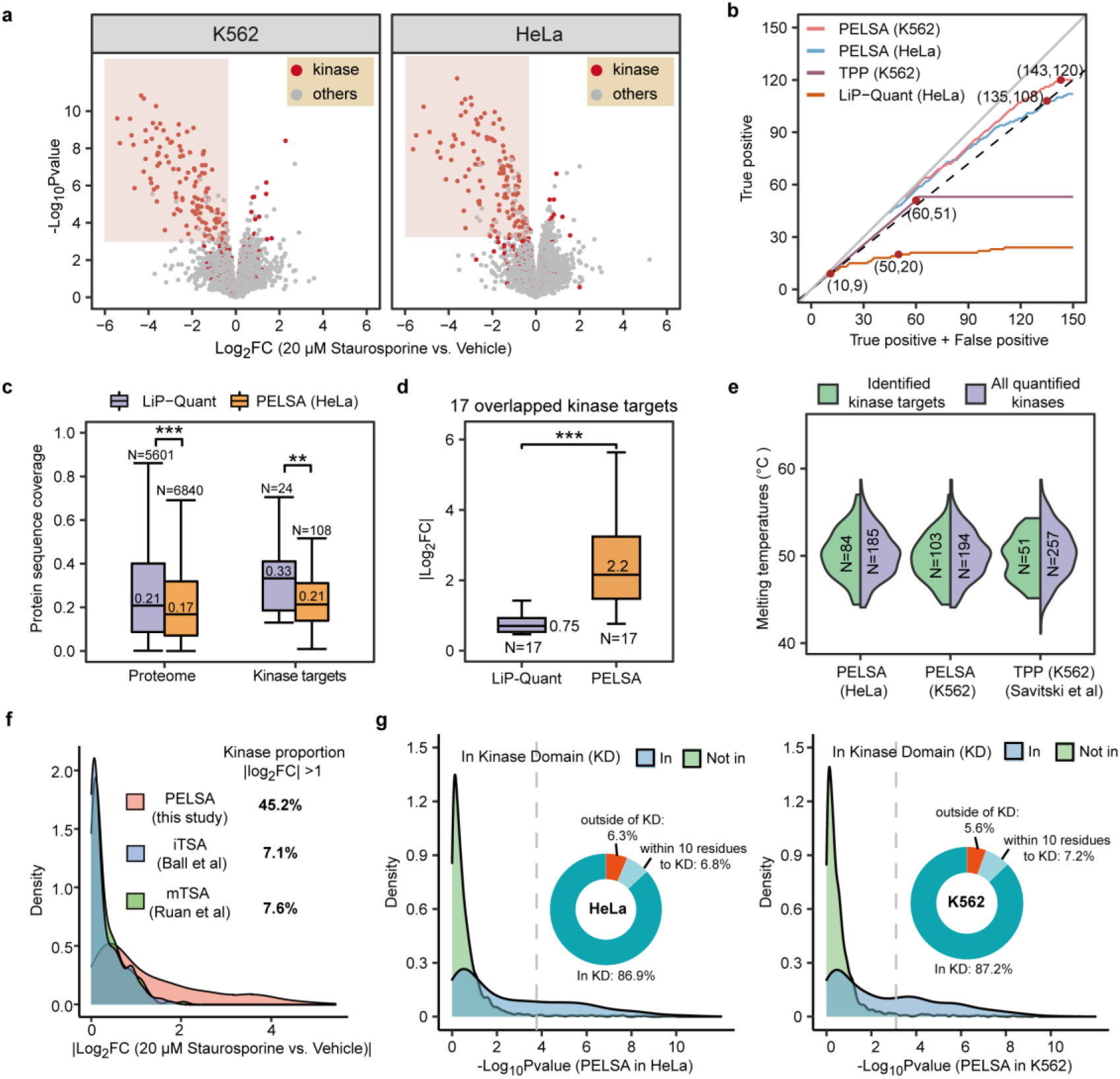
Comparing target identification performance of PELSA with existing modification-free methods. **a,** Volcano plot visualization of all proteins from PELSA analyses of K562 (left) and HeLa (right) lysates exposed to 20 μM staurosporine. The lower boundary of the red shadow denotes the threshold of -log_10_Pvalue, above which over 80% of the stabilized proteins (log_2_FC < 0) are kinases. **b,** True positive rate (TPR) evaluation for the selected assays in staurosporine target identification. The labeled points represent the numbers of identified candidate targets and kinase targets in each assay (TPR up to 80%). LiP-Quant is also labeled at the kinase target number of 20 (TPR = 40%). The grey line (slope = 1) and black dashed line (slope = 0.8) represent 100% and 80% of the candidate targets are kinase targets, respectively. **c, d,** **p < 0.01 and ***p < 0.001, Wilcoxon signed-rank test; medians are labeled and other settings are as Fig.1b. (**c**), Protein sequence coverages for the whole quantified proteome (left) and identified kinase targets (right) in LiP-Quant HeLa and PELSA HeLa analyses. (**d**), Fold changes of kinase targets that were identified by both LiP-Quant (using TPR cutoff of 40%) and PELSA (HeLa). **e,** Comparing melting temperatures (Tm) of identified kinase targets and all quantified kinases in the TPP dataset and two PELSA datasets. Some PELSA kinase targets lack TPP-reported Tm values. **f,** Comparing the fold changes of kinases quantified in PELSA, iTSA, and mTSA. **g,** Density plots showing -log_10_Pvalue distributions of peptides with tryptic cleavage sites located in and outside the kinase domains for K562 and HeLa PELSA analyses. The dashed lines indicate the significance cutoffs defined in (**a**). The doughnut charts show the location distributions of the kinase peptides that passed the significance cutoffs. *Note*: Kinase targets refer to kinase proteins that are identified as staurosporine-binding proteins; quantified kinases refer to all kinases in the dataset including kinase proteins that are not identified as staurosporine-binding proteins. LiP-Quant, TPP, iTSA, and mTSA datasets were retrieved from the literatures^10, 12, 25, 27^.

PELSA with one E/S ratio also identified 2.4-fold more kinase targets than TPP using ten temperatures (120 versus 51) in lysates of the same cell line (K562) (**Fig. 2b**), although more proteins were included in the TPP dataset (7638 versus 6310)^10^. TPP showed a bias against the thermo-resistant and thermo-susceptible kinases, whereas PELSA is capable of identifying kinase targets with extreme melting temperatures (**Fig. 2e**). Moreover, PELSA also substantially outperforms the recently updated versions of TPP—iTSA, 2D-TPP, and mTSA—for staurosporine target identification: compared to PELSA’s 120 kinases/143 candidates, iTSA identified 71 kinases/85 candidates^25^; 2D-TPP identified 60 kinases/73 candidates^26^ and mTSA identified 64 kinases/85 candidates^27^ (**Supplementary Table 1**). We compared the readouts of kinases in PELSA and in iTSA and mTSA, which also determine target proteins via output of abundance fold changes of proteins (*i.e*., readout). The comparison results showed that 44.5% of the kinases in the PELSA staurosporine dataset displayed readout of >2, while the proportions were 7.1% and 7.6% in iTSA staurosporine and mTSA staurosporine datasets, respectively (**Fig. 2f**). Beyond the high sensitivity in target protein identification, PELSA also brings along the capacity to identify binding regions: for kinase targets identified by PELSA, over 93% of the peptides passing the significance cutoff (**Supplementary Discussion**) are located in or within 10 residues away from the known staurosporine-binding domain—kinase domain (**Fig. 2g**).

### Exploring weak metabolite-protein interactions

Encouraged by the excellent performance of PELSA in drug target identification, we next examined whether PELSA is capable of detecting weak metabolite-protein interactions by investigating the binding proteins of two metabolites—folate and leucine—which are known to bind their target proteins with micromole-level affinity^18, 28, 29^.

Folate PELSA analysis successfully identified dihydrofolate reductase DHFR (a known folate-binding protein)^18^ as the top hit (**Fig. 3a**) and revealed that the top five DHFR peptides with the most profound stabilization were mainly present in the folate-binding pocket (**Fig. 3b**). Beyond DHFR, PELSA also identified three Uniprot-annotated folate-analog-binding proteins, *i.e*., MTHFR, GART, and ATIC among the top 6 most significantly stabilized proteins by folate treatment (**Fig. 3a**); PELSA revealed that they were all stabilized at known folate-analog-binding sites (**Fig. 3c-e**), indicating that folate may compete with these analogies to bind their cellular targets. The 3rd most significantly stabilized protein was a collagen proline hydroxylase, P3H1 (**Fig. 3a**). A previous report indicates that folate may function as a reducing agent to participate in the hydroxylation of collagen proline^30^. Coincidentally, the local stability profiles revealed that, albeit with 11 peptides of P3H1 quantified, only the three peptides from the prolyl 4-hydroxylase domain were stabilized by folate treatment (**Fig. 3f**). Our results thus provide evidence for the participation of folate in the hydroxylation of collagen proline.

**Fig. 3.**
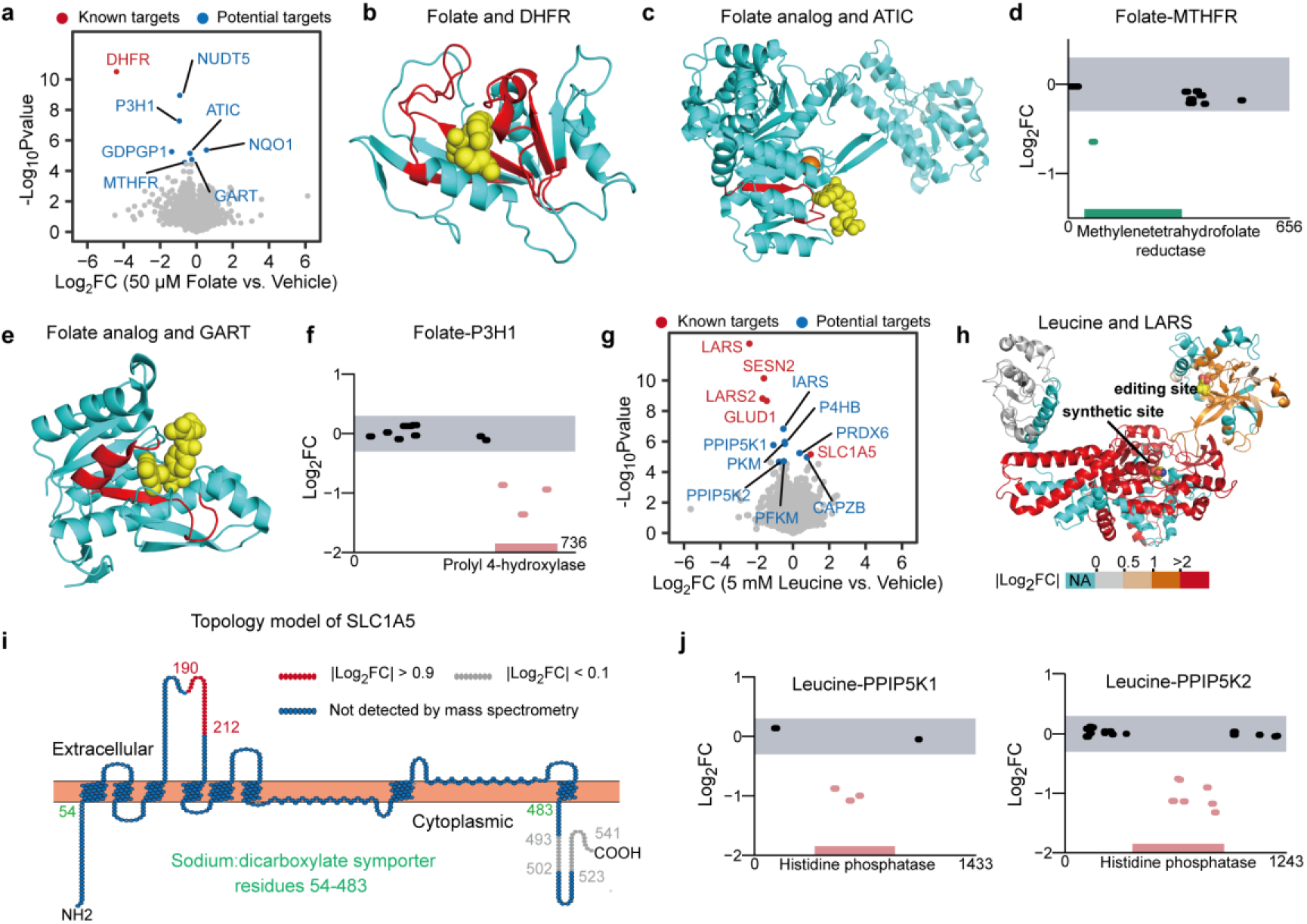
Detecting weak metabolite-protein interactions. **a**, Volcano plot visualization of all proteins from a PELSA analysis of K562 lysates exposed to 50 μM folate. **b,** Complex structure of folate and DHFR generated by superposition of human DHFR (PDB: 1BOZ) against *E.coli* DHFR-folate complex (PDB: 4EJ1). Folate, yellow spheres; the top five peptides with the most profound stabilization (Bayes t-test, -log_10_Pvalue > 2, ranked by -log_2_FC) are colored in red. **c,** Complex structure of folate analog (yellow spheres) and ATIC (PDB: 1P4R). The peptide with most profound stabilization is colored in red. **d,** Local stability profiles of MTHFR by 50 μM folate treatment. **e,** Complex structure of folate analog (yellow spheres) and GART (PDB: 1RBY). The absolute log_2_FC values of all quantified GART peptides are < 0.5, and thus the top two peptides with the lowest p value (Bayes t-test) are colored in red. **f,** Local stability profiles of P3H1 by 50 μM folate treatment. **g,** Volcano plot as in (**a**) but of analysis of K562 lysates exposed to 5 mM leucine. **h,** Structure of LARS in complex of leucine (multicolor spheres) both in the editing site and the synthetic site (PDB: 6KQY) with peptides colored based on their log_2_FC values. **i,** Topology model of SLC1A5 generated by Protter^64^. Protein sequences are colored based on their log_2_FC values. **j,** Local stability profiles of PPIP5K1 (left) and PPIP5K2 (right) by 5 mM leucine treatment.

The top four hits (LARS, SESN2, LARS2, and GLUD1) identified for leucine were all well-known leucine-binding proteins (**Fig. 3g**). Notably, LARS contains two leucine-binding sites^28^: the synthetic site and the editing site. Our PELSA data revealed that both of the leucine-binding sites were stabilized upon leucine binding, but with distinct magnitudes (**Fig. 3h**). This observation supports the potential of PELSA to informatively differentiate between discrete ligand-binding sites in a single protein.

Our PELSA data also showed that leucine treatment destabilized SLC1A5 (**Fig. 3g**), an amino acid transporter located at the plasma membrane that can accept leucine as a substrate^31^. While the PELSA dataset included three SLC1A5 peptides, only one was destabilized (**Fig. 3i**): the peptide located at the extracellular segment of the substrate-binding domain (residues 54 to 483)^32^, suggesting that leucine binding may induce this segment to adopt a more flexible conformation. Beyond known leucine-binding targets, PELSA identified additional putative leucine targets including PPIP5K1 and PPIP5K2 (**Fig. 3g**), which are reported to involve with cancer cell proliferation^33, 34^. Notably, PELSA revealed that leucine binds both PPIP5K1 and PPIP5K2 at the conserved functional histidine phosphatase domains (**Fig. 3j**), which may provide clues for future function studies of leucine.

### Characterizing the recognition domains of PTMs

Post-translational-modifications (PTMs) can be recognized by downstream effector proteins (so-called ‘‘readers’’) through the recognition domains (**Fig. 4a**) to regulate cellular events^5^. However, the interactions between PTMs and reader proteins are often weak and transient. Despite recent progress in modification-based methods^5, 35^, it remains challenging to identify reader proteins and recognition domains of PTMs in complex cellular environment. We wondered whether PELSA is able to fill this gap.

**Fig. 4.**
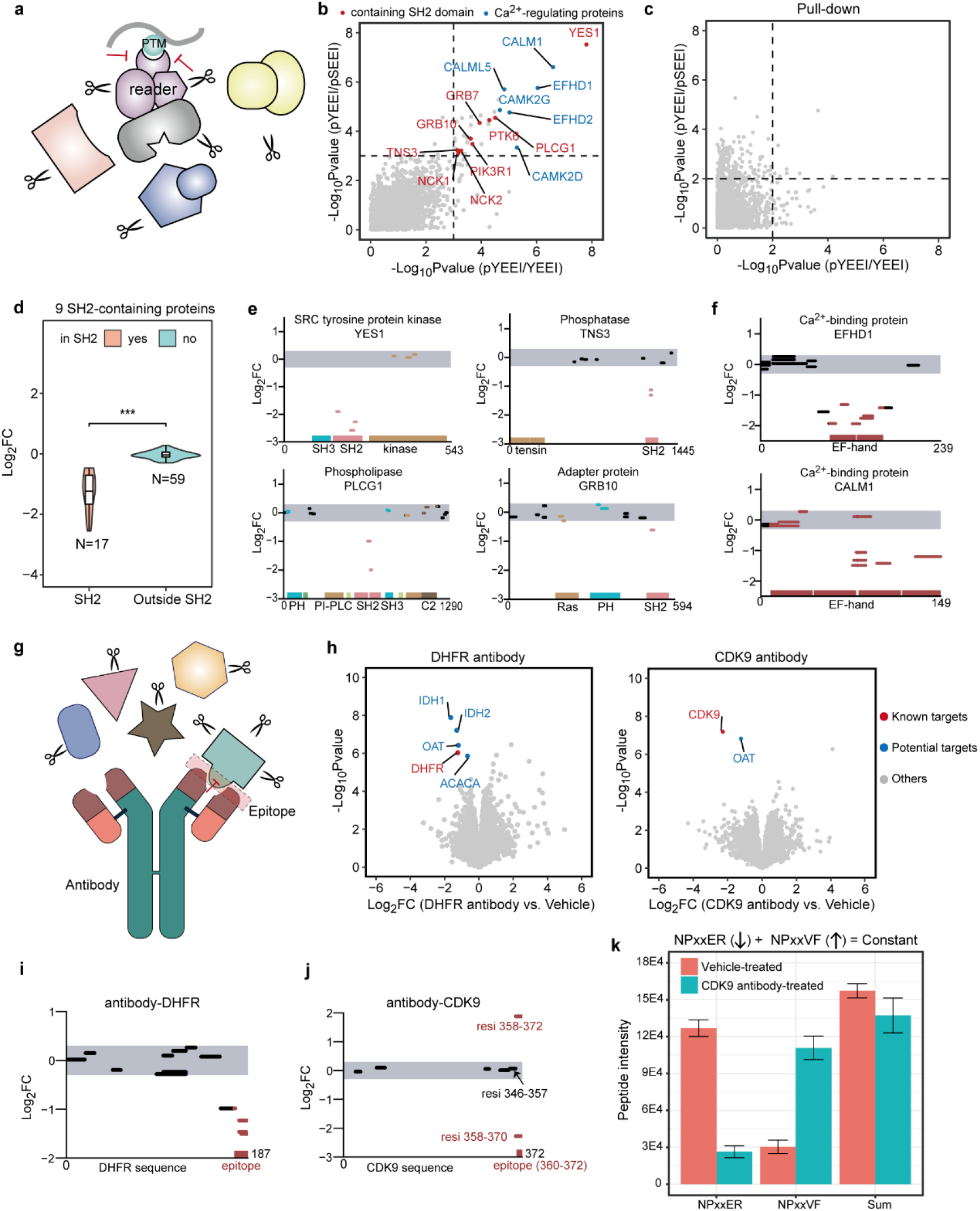
Identifying recognition domains of a PTM and localizing epitopes of antigens. **a,** Schematic representation of PELSA to reveal the PTM-recognition domain. **b,** Scatter plot of protein -log_10_Pvalues in PELSA (pYEEI/YEEI) and PELSA (pYEEI/pSEEI) (**Methods**). The dashed lines indicate the significance cutoff (-log_10_Pvalues = 3.1). The proteins passing the significance cutoff are colored: SH2-containing proteins, red; Ca^2+^-regulating proteins, blue; others, grey. **c,** Scatter plot as in (**b)** but for the pulldown experiment (three lysate replicates). The dashed lines indicate a relaxed significance cutoff (-log_10_Pvalues = 2). No SH2-domain containing proteins passed the significance cutoff. **d**, Log_2_FC distributions of the peptides (from 9 SH2-containing target proteins) that reside in and out of the SH2 domains. Violin plots represent relative densities and the settings of the inner boxplots are as Fig. 1b. ***p < 0.001, Wilcoxon signed-rank test. **e and f,** Local stability profiles of SH2-containing proteins from different protein families (**e**) and representative Ca^2+^-regulating proteins (**f**) by pYEEI treatment. **g,** Schematic representation of PELSA to reveal the epitope. **h**, Volcano plot visualization of all proteins from PELSA analyses of HeLa lysates exposed to DHFR antibody (left) or CDK9 antibody (right). **i** and **j**, Local stability profiles of DHFR and CDK9 by DHFR antibody and CDK9 antibody treatment, respectively. **k**, Intensities of NPxxVF and NPxxER, and their sum. Four replicates (mean ±S.D.).

Here we focused on phosphotyrosine (pY), exemplified by the pYEEI motif which preferentially binds Src-kinase SH2 domains^36^. PELSA revealed that 28 proteins were significantly stabilized by pYEEI, among which 9 proteins contain SH2 domains (**Fig. 4b**). By contrast, we did not identify any SH2-domain-containing proteins in our pulldown experiment (**Fig. 4c**), possibly because the weak interactions between pY and its reader proteins^37^ are susceptible to loss during the stringent washing procedure. Beyond the advantages over pulldown in detecting weak PTM-protein interactions, PELSA also featured with recognition domain identification. As anticipated, peptides located in the SH2 domains displayed a significantly reduced abundance, whereas peptides out of the SH2 domains remained unchanged (**Fig. 4d**). Notably, SH2 domains from different protein families were stabilized by pYEEI with varying magnitudes (**Fig. 4e** and **Supplementary Table 2**): in accordance with the binding preference of pYEEI^36^, the SH2 domain of the Src kinase YES1 displayed the most profound stabilization. Several Ca^2+^-regulating proteins were found stabilized by pYEEI at the Ca^2+^-regulating regions (**Fig. 4f** and **Supplementary Table 2**), although the underlying mechanism is unclear.

Although only recognition domains of pY were investigated here, it is reasonable to further extend the application scope to investigate the recognition domains of many other PTMs, which is crucial to understand the biological functions of the PTMs.

### The high-resolution binding data of PELSA enables epitope identification

We then asked whether PELSA can determine the ligand-binding regions when the ligand is a protein such as an antibody (**Fig. 4g**). To this end, two commercial antibodies (against DHFR or CDK9) were investigated with PELSA using HeLa cell lysates. PELSA quantified 6806 and 6207 proteins in DHFR and CDK9 antibody experiments, respectively, and the corresponding antigen proteins DHFR and CDK9 were found in the top 5 most significantly stabilized proteins in respective experiments (**Fig. 4h**). Of note, most of the significantly stabilized non-antigen proteins (-log_10_Pvalue > 5, log_2_FC < 0, Bayes t-test) contain multiple stabilized peptides (**Supplementary Table 3**) indicating the high confidence of their interactions with the added antibody, possibly resulting from the low specificity of the antibodies.

Four out of the 15 quantified DHFR peptides displayed a significantly reduced abundance (Bayes t-test p < 0.01, log_2_FC < -0.3) upon DHFR antibody binding (**Fig. 4i**). Strikingly, their tryptic cleavage sites were located exactly in the known epitope (residues 172-187) (**Fig. 4i**). CDK9 antibody recognizes a 13-amino-acid epitope— sequence PATTNQTEFERVF (residues 360-372)—which is located at the tail of CDK9. The local stability profiles revealed that peptide NPATTNQTEFER (NPxxER, residues 358-370) with C terminus cleavage site located exactly in the epitope displayed a significantly reduced abundance (Bayes t-test p < 0.001, log_2_FC = -2.28) (**Fig. 4j**), whereas even the peptide (residues 346-357) with C terminus two residues away from the epitope remained unchanged (-log_10_Pvalue = 0.153, log_2_FC = 0.07, Bayes t-test). Notably, the missed cleavage form of NPxxER—NPxxVF (residues 358-372)—displayed an opposite direction of change with NPxxER (**Fig. 4j**), which can be explained: binding with antibody inhibited the trypsinization at residue 370; thus, less amount of NPxxER (residues 358-370) was generated, and thereby more NPxxVF (residues 358-372) was left (**Fig. 4k**). Overall, these results indicated that PELSA can identify the epitopes on antigen proteins at high resolution.

### Assaying Zn^2+^ responsive regions across the proteome

Next, we wondered whether PELSA is applicable for the ligand with a small size, like a single-atom metal ion—Zn^2+^, which typically binds protein on a small zinc-finger (ZnF) motif composed of ∼30 amino acids^38^. The cell lysates depleted of endogenous Zn^2+^ were treated with varying concentrations of Zn^2+^ or vehicle, and then subjected to PELSA analysis (**Extended Data Fig. 5a**). After 30 μM Zn^2+^ treatment, 280 proteins were significantly stabilized (-log_10_Pvalue > 3, log_2_FC < -0.5, Bayes t-test), among which ∼68% (190 proteins) were Uniprot-annotated metal-binding proteins (**Fig. 5a,b**). This proportion was substantially higher than that in the measured proteome (19%) (**Fig. 5b**). Among the 190 metal-binding proteins identified by PELSA, 112 are Uniprot-annotated Zn^2+^-binding proteins, and 78 are proteins known to be bound by other divalent metal ions highlighted by Ca^2+^, Mg^2+^, Fe^2+^, and Mn^2+^ (**Fig. 5c** **and Supplementary Table 4**), indicating a divalent-metal-ion promiscuity, which is frequently observed in metal-binding proteins^39^. Zn^2+^-binding proteins have been previously investigated with chemical probes^40^. In that study, 38 putative Zn^2+^-binding proteins were determined: 6 were Uniprot-annotated Zn^2+^-binding proteins and 9 were other-metal-binding proteins (**Supplementary Table 5**). This comparison demonstrated the excellent performance of PELSA in identifying metal-binding proteins.

**Fig. 5.**
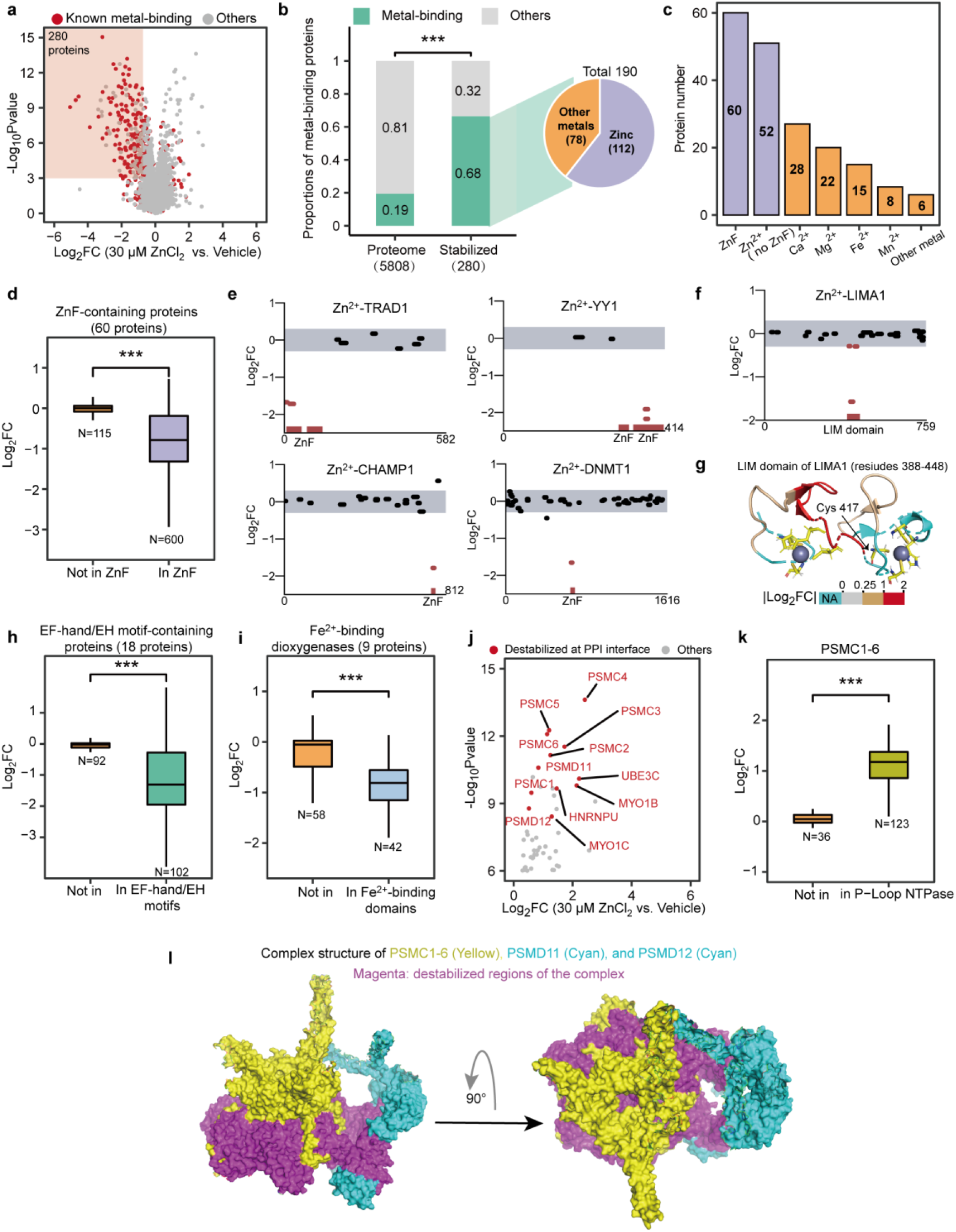
Characterization of Zn^2+^ proteome revealing the stabilized metal-binding regions and destabilized protein-protein interfaces. **a,** Volcano plot visualization of all proteins from a PELSA analysis of HeLa lysates exposed to 30 μM ZnCl_2_. The right boundary and lower boundary of the red shadow denote log_2_FC of -0.5 and - log_10_Pvalue of 3, respectively. **b,** Proportions of metal-binding proteins in the whole dataset and in the significantly stabilized subset. ***p < 0.001, Fisher’s exact test. Pie-chart denotes the percentage of known Zn^2+^-binding proteins among all the stabilized metal-binding proteins. **c,** Compositions of the metal-binding proteins that were stabilized by 30 μM Zn^2+^ treatment. **d, h, i, and k,** Log_2_FC distributions of peptides that reside in and out of the indicated domains. ***p < 0.001, Wilcoxon signed-rank test. (**h**), EF-hand/EH motifs are known Ca^2+^-binding motifs^65^. (**k**), P-loop-NTPase domains are the binding surfaces of the adjacent members of PSMC complex^66^. **e,** Local stability profiles of representative ZnF-containing proteins. **f,** Local stability profiles of LIMA1. **g,** LIM domain of LIMA1 (PDB: 2D8Y) with peptides colored based on log_2_FC values. Zn^2+^-binding residues: yellow sticks; zinc ions: dark-purple spheres. **j,** The zoom-in view of the volcano plot that displayed in **(a)**. **l,** Surface representation of PSMC1-6, PSMD11, and PSMD12 complex (PDB: 5LN3) viewed from the lateral side with PSMC3 exposed (left) and viewed from the top (right). This complex is destabilized at the interacting surfaces of its members (colored in magenta).

Beyond precisely determining the small Zn^2+^-binding sites (**Fig. 5d-g, Supplementary Discussion**), PELSA revealed that Zn^2+^ stabilized the Ca^2+^-, Fe^2+^-, and Mg^2+^-binding proteins at Ca^2+^-, Fe^2+^-, and Mg^2+^-binding regions, respectively (**Fig. 5h,i, Extended Data Fig. 5b, and Supplementary Table 6)** which is agreement with previous findings that Zn^2+^ can occupy the binding pockets of other divalent metal ions^41, 42^. Our PELSA analysis also provided a local stability atlas of 90 Zn^2+^-stabilized proteins that were not categorized as metal binding (**Extended Data Fig. 5c**); Gene ontology analysis of these 90 proteins revealed an enrichment of GTP-binding proteins (**Extended Data Fig. 5d**), particularly Ras-related proteins (**Supplementary Table 4**). One Ras-related protein, RAB1A, has been previously reported as a Zn^2+^-buffering protein^43^. Our results indicate a potential prevalent role of Ras-related proteins in regulating cellular Zn^2+^ homeostasis.

Proteins will be destabilized, if ligands bind to their partner proteins and dissociate the partner proteins from the formed protein complexes^10, 44^. Among the top 18 proteins destabilized by Zn^2+^ (log_2_FC > 0, ranked by -log_10_Pvalue, Bayes t-test), 12 proteins were destabilized at known protein-protein interaction interfaces (**Fig. 5j-l, Extended Data Fig. 5e**, **Supplementary Table 6**, **and Supplementary Discussion**). This specific destabilization is also recapitulated in PELSA 20 μM Zn^2+^ analysis (**Extended Data Fig. 5f,g and Supplementary Table 6)**, suggesting the potential of PELSA to monitor the assembly states of protein complexes.

### Target landscapes of α-ketoglutarate and R-2-hydroxyglutarate in HeLa and Jurkat cells

Isocitrate dehydrogenase (IDH) gene mutations are frequently observed in multiple human cancers^45^; these mutations can impart a neomorphic enzyme activity wherein α-ketoglutarate (αKG) can be converted to the R enantiomer of 2-hydroxyglutarate (R2HG)^46^. R2HG is structurally similar to αKG (**Fig. 6a**) and has been reported to act as a weak competitive inhibitor of multiple αKG-dependent dioxygenases (KGDDs)^47^. The highly simple and similar structures of these two metabolites make it challenging to identify their binding proteins through modification-based methods. Moreover, the low affinity of these two metabolites, especially R2HG (often up to millimole-level affinity)^48^, further exacerbates the difficulty of target identification. As a result, despite wide-recognized roles of R2HG in cancer development^49, 50^, there is no proteome-wide investigation of R2HG binding proteins.

**Fig. 6.**
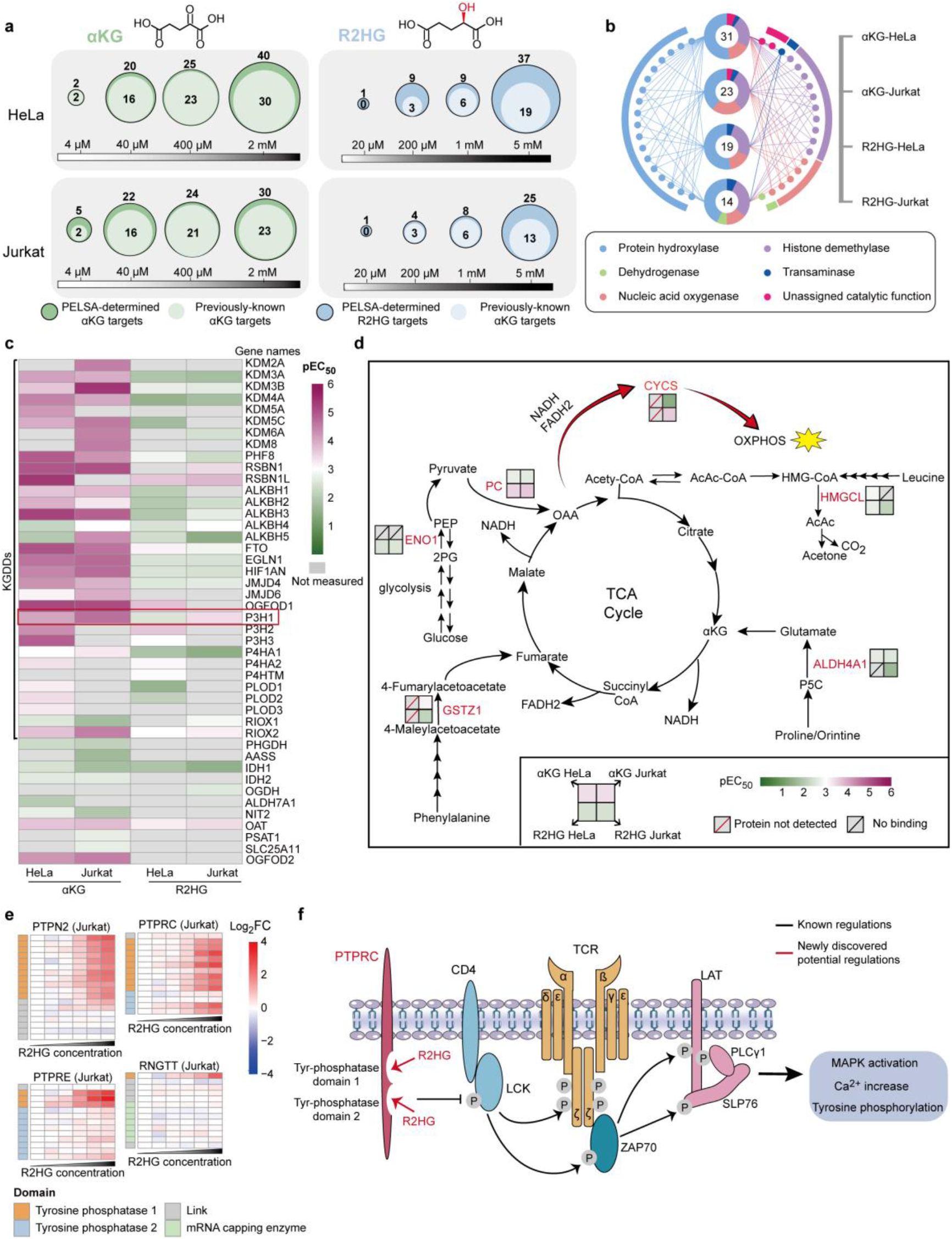
Characterizing binding profiles of αKG and R2HG in two cell lines. **a,** Bubble plots displaying the numbers of αKG and R2HG targets identified by PELSA (**Supplementary Discussion**). The inner bubble denotes previously-known αKG targets; the outer denotes all candidate αKG/R2HG targets identified by PELSA. **b,** The radiation diagram depicts categories of the previously-known αKG targets identified in αKG-HeLa, αKG-Jurkat, R2HG-HeLa, and R2HG-Jurkat. The central donut reflects the proportions of each protein category occupied in the annotated PELSA analysis. Each node around the cycle denotes one protein. The linkage between the node and the donut denotes the protein is identified as a target protein in this PELSA analysis. The labeled number denotes the count of previously-known αKG targets identified in each analysis (total count identified across all concentrations). **c,** Heatmap displaying pEC_50_ values of 44 previously-known αKG targets toward αKG and R2HG (measured by PELSA in both HeLa and Jurkat cell lysates). Grey cells in the heatmaps indicate no measurements. **d**, Schematics of simplified glycolysis, TCA cycle, amino acid metabolism, and OXPHOS pathways. The putative αKG and R2HG targets are marked in red with binding affinities indicated. **e,** Local affinity profiles of four tyrosine-phosphatase-domain-containing proteins for R2HG treatment in Jurkat cell lysates. **f,** PTPRC is an upstream regulator of TCR signaling. R2HG destabilized PTPRC at its functional domains.

We used PELSA to explore the binding proteins of αKG and R2HG in lysates of HeLa and Jurkat cells. PELSA analysis of 2 mM αKG treated HeLa cell lysates identified 40 significantly stabilized proteins (-log_10_Pvalue > 3.4, log_2_FC < -0.5, Bayes t-test), among which 30 are previously-known αKG targets (65 in total in this dataset; **Extended Data Fig. 6a,b**)^51^. This represents the largest number of known αKG targets identified in a single analysis. Although αKG has been investigated in a LiP-MS study with *E.coli* lysates^52^, only 2 previously-known αKG targets were identified (33 in total in the LiP-MS dataset; **Extended Data Fig. 6b**).

As anticipated, PELSA R2HG analyses identified fewer known αKG targets than PELSA αKG analyses in both HeLa and Jurkat cell lysates (**Fig. 6a**). The cell-line comparison revealed that protein-hydroxylase targets were underrepresented in Jurkat cells (**Fig. 6b**) relative to HeLa cells, which can be explained by the differential expression levels of protein hydroxylases in these two cell lines (**Extended Data Fig. 6c**).

PELSA also enables the determination of αKG-binding regions for tens of αKG targets in a single analysis (**Extended Data Fig. 6d and Supplementary Table 7**). Previous co-crystal structural studies of purified KDM4A in complex with R2HG revealed that R2HG occupies the same binding pocket as αKG^47^. Note that for multiple αKG targets, our PELSA data of both HeLa and Jurkat cell lysates indicate that R2HG binds the same pockets as αKG (**Extended Data Fig. 6d and Supplementary Table 7**).

PELSA determined the binding affinities between αKG (R2HG) and 44 previously-known αKG targets in lysates of HeLa and Jurkat cells (**Fig. 6c**). In agreement with previous findings^47^, PELSA revealed that R2HG has lower binding affinities for KGDDs compared to αKG (**Fig. 6c**). Although the binding affinities in the two cell lysates are well correlated (**Extended Data Fig. 6e)**, we observed that P3H1 displayed a higher affinity for both αKG and R2HG in Jurkat cell lysates compared to HeLa cell lysates (**Fig. 6c** **and Extended Data Fig. 6f**), which may represent the distinct regulating factors (*e.g*., interacting partners and post-translational modifications) of P3H1 in HeLa and Jurkat cells.

### Previously unknown targets of αKG and R2HG

PELSA identified 19 high-confidence (**Supplementary Discussion**) previously-unknown targets of αKG or R2HG and determined their binding affinities in both HeLa and Jurkat cell lysates (**Supplementary Table 8**). Notably, many of these proteins are involved with energy metabolism, including amino acid metabolism, glycolysis, oxidative phosphorylation (OXPHOS), and TCA cycle anaplerosis (**Fig. 6d**). Interestingly, different from KGDDs which bind more strongly to αKG than R2HG, pyruvate carboxylase (PC), an enzyme critical for TCA anaplerosis, was identified to bind both αKG and R2HG, but with higher affinity toward R2HG than αKG in lysates of both HeLa and Jurkat cells (**Fig. 6d** and **Extended Data Fig. 7a**). This finding was also confirmed by dose-response experiments via western-blot readouts (**Extended Data Fig. 7b**). The PELSA local affinity data also revealed that R2HG stabilized PC on the segment responsible for the transfer of carboxy group to pyruvate (**Extended Data Fig. 7a**)^53^. We purified this segment and verified the stabilization by R2HG using a thermal shift assay (**Extended Data Fig. 7c**). IDH mutations can lead to remarkably high R2HG levels, accompanied by disruption of redox homeostasis and alteration of amino acid metabolism and TCA cycle anaplerosis^46^. Little is known about whether R2HG has a role in these metabolism alterations and how R2HG functions. Our PELSA evidence for the interactions between R2HG and the proteins (involved with the TCA cycle anaplerosis, amino acid metabolism, and OXPHOS) (**Fig. 6d**) thus yields an insight into the aberrant cellular metabolism in IDH-mutated cancer cells.

Beyond the enzymes with well-known functions, we also identified two putative enzymes without known substrates, *i.e.,* HDHD2 and FAHD2A (**Supplementary Table 8**); their interactions with αKG/R2HG and relative binding affinities to these two metabolites (**Extended Data Fig. 7d)** may afford clues for their biological functions.

In addition to the targets stabilized by αKG and R2HG, we found that a group of tyrosine-protein phosphatase domain-containing proteins—PRPRC, PTPRE, PTPN2, and RNGTT—were destabilized exclusively in R2HG-treated Jurkat cell lysates (**Extended Data Fig. 8a,b**). Moreover, these proteins were all destabilized at their shared tyr-protein phosphatase domains (**Fig. 6e**); this R2HG-induced Jurkat-specific destabilization was also confirmed by another biological replicate of PELSA R2HG analysis (**Extended Data Fig. 8c,d**). R2HG has been reported to suppress T cell receptor (TCR) signaling^54^. Notably, PTPRC, a membrane protein that functions as a gatekeeper of TCR signaling^55^, was also among the R2HG-destabilized proteins. PTPRC is known to employ its tyr-protein phosphatase domains to regulate TCR signaling (by dephosphorylating, and thus activating LCK) (**Fig. 6f**). Our PELSA data indicating that R2HG destabilized PTPRC’s tyr-protein phosphatase domains therefore uncovers a possible basis to help explain previous reports of R2HG-mediated suppression of TCR signaling^54^. Overall, our dose response PELSA analyses of αKG and R2HG in the two cell lines provide informative interaction data for future hypothesis generation studies of αKG and R2HG.

## DISCUSSION

In this study, we found that disruptive trypsinization amplifies the readout of ligand-induced protein local stability shifts, and developed this concept into a powerful technology—PELSA—which allows simultaneous sensitive target protein identification and ligand-binding region determination in native cellular environment without ligand modification. Compared against existing modification-free methods that enable binding region determination (LiP-MS methods)^12^, PELSA identified 6-fold more FKBP family target proteins (6 versus 1) for rapamycin and 12-fold more kinase targets (108 versus 9) for a pan-kinase inhibitor (staurosporine) than LiP-MS and LiP-Quant, respectively. Compared with prevalent modification-free methods that do not yield binding region information (TPP methods), PELSA identified 1.7-2.4 times more kinase targets for staurosporine than TPP and recently revised TPP methods (iTSA, 2D-TPP, and mTSA).

Beyond high sensitivity in target identification, PELSA’s peptide-level readout also enables binding-region determination. PELSA detects the ligand-induced local stability shifts to deduce ligand-binding regions. In some cases, *i.e*., when the binding signals are propagated to distal locations within the domain through cooperative intra-segment interactions^56, 57^, PELSA can accurately and sensitively determine the ligand-binding domains. For example, PELSA simultaneously determined staurosporine-binding domains for 120 kinases, which represents the largest number of ligand-binding regions determined in a single analysis. In other cases, *i.e.*, when ligand binding only affects the stability of certain residues of the proteins, PELSA can determine the binding residues. This was demonstrated by determining a 13-amino-acid epitope for an antibody and Zn^2+^ binding residues within a 60-amino-acid domain.

Our study also provides a powerful solution for identifying recognition domains of PTMs. A recent study reported that a tri-functional amino acid can enable identifying PTM-binding regions when it is placed 1 or 2 residues away from the PTM sites of interest^5^. However, the case of phosphotyrosine (pY) binding has shown that alteration of the +2 and +3 positions can profoundly alter the binding profiles of pY^58^. PELSA does not require prior modification of the analyte ligand, and we have successfully applied PELSA to characterize the recognition domains of pY in this study. Given the ubiquity of PTM-mediated regulation in biology and the many pathological associations of dysregulation PTMs^59, 60^, PELSA’s ability to identify recognition domains of PTMs in human cell lysates will almost certainly motivate its use in many, highly diverse biological and medical studies.

We also showcase the capacity of PELSA for sensitively and informatively probing weak interactions by identifying the binding proteins of leucine, folate, αKG, and R2HG. While previous studies have employed modification-based or modification-free methods to investigate metabolite-binding proteins^52, 61, 62^, these approaches often generate a large number of candidate targets with a limited number of known metabolite-binding proteins. In contrast, PELSA results consistently exhibit a significantly higher percentage of known-binding events. For instance, in a prior LiP-MS study of αKG-treated *E.coli* lysates^52^, 34 candidate targets were identified with 2 known αKG binding proteins. In comparison, PELSA identified 40 candidate targets, and notably, 30 of these were known αKG binding proteins, despite using a more complex lysate sample (human HeLa cell lysate). We envision that PELSA’s improved hit rate has the potential to significantly streamline the validation process in hypothesis-generation studies.

In summary, we demonstrate PELSA is a highly sensitive and generic method to reveal binding regions on proteins of very diverse ligand types (including drugs, antibodies, phosphorylated peptides, metal ions, and metabolites) on a proteomics scale, without the need for chemical modification of the analyte ligand. Beyond ligand binding, the transition of a protein between different proteoforms (*e.g*, the presence or absence of post-translational modification)^63^ may also induce protein stability shifts, and thus could also be investigated by PELSA. We envision that PELSA will find wide utilization throughout life science research.

## ACKNOWLEDGMENTS

This work was supported, in part, by funds from the National Key Research and Development Program of China (2021YFA1302601, 2020YFE0202200 to M.Y.), the National Natural Science Foundation of China (92153302, 22034007 to M.Y., 91853205, 81625022 to C.L., and 22137002 to K.W.), the innovation program of science and research from the DICP, CAS (DICP I201935, DICP I202139 to M.Y.), the project of National Multidisciplinary Innovation Team of Traditional Chinese Medicine from National Administration of Traditional Chinese Medicine (ZYYCXTD-202004 to C.L.), the Youth Innovation Promotion Association of CAS (No.2022279 to S.C.), and Supported by Innovation Academy for Precision Measurement Science and Technology, CAS to M.Y.

## AUTHOR CONTRIBUTIONS

M.Y., K.L., and K.W. conceived and designed the project; K.L. developed the method, carried out the experiments, and analyzed the results under the supervision of M.Y.; S.C., F.Y., and S.G. measured the binding affinities of purified HSP90AA1 to HSP90 inhibitors and validated the off-targets identified for HSP90 inhibitors and R2HG under the supervision of C.L.; K.W. advised on the experimental design. Y.W. set up the MS analysis methods, discussed the experiment results, and edited the manuscript. Z.F. gave help with data processing. J.L. helped in the experiment of investigating PTM’s readers. H.Z. helped in data processing in the Zn^2+^ experiment. Y.L. helped to correct the manuscript; T.Y. advised on investigating the leucine-binding proteins. J.Z. helped in calculating the Euclidean distances between peptides and ligands. X.Z., C.R., and Q.W. gave scientific advice; K.L. and M.Y. wrote the paper with input from other authors.

## Data and materials availability

The raw mass spectrometry proteomics data, protein identification and quantification results have been deposited with the ProteomeXchange Consortium via the PRIDE partner repository with the dataset identifier PXD034606 and will be made accessible upon publication.

---

**Extended Data Fig. 1.**
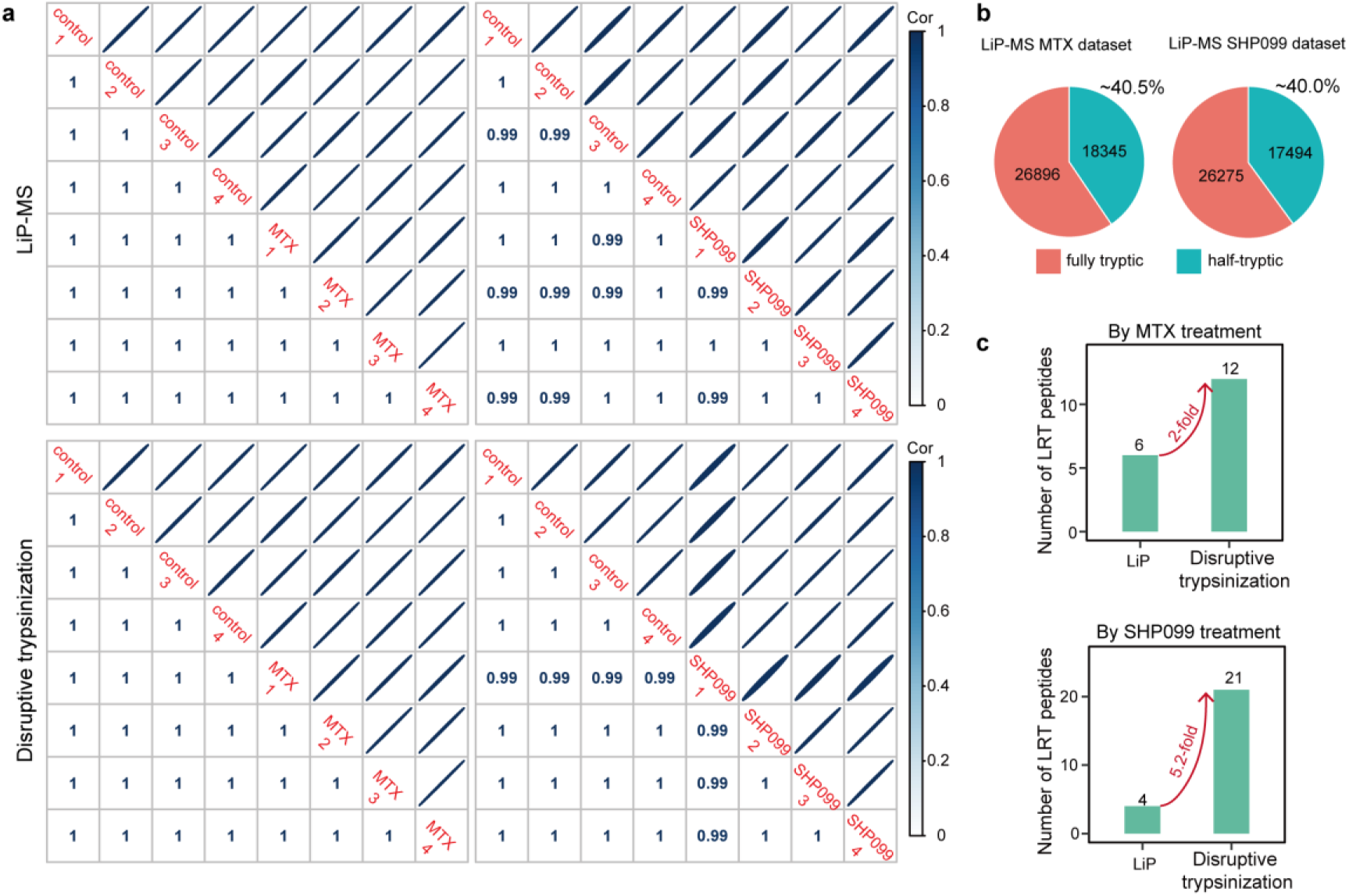
Data quality assessment of in-house performed LiP-MS experiments. **a**, Intensities of peptides generated by LiP-MS (top) or disruptive trypsinization (bottom) show excellent correlations across replicates. **b**, The proportions of half-tryptic peptides in our in-house-performed LiP-MS experiments (40.5% and 39.9%), agree well with that reported in the literature (*i.e.*, 40%)^11^. **c**, Bar-plots displaying the numbers of ligand-responsive target (LRT) peptides (*i.e.*, target protein peptides that showed |log_2_FC| > 0.3 & -log_10_Pvalue > 2) in the LiP-MS datasets and disruptive trypsinization datasets.

**Extended Data Fig. 2.**
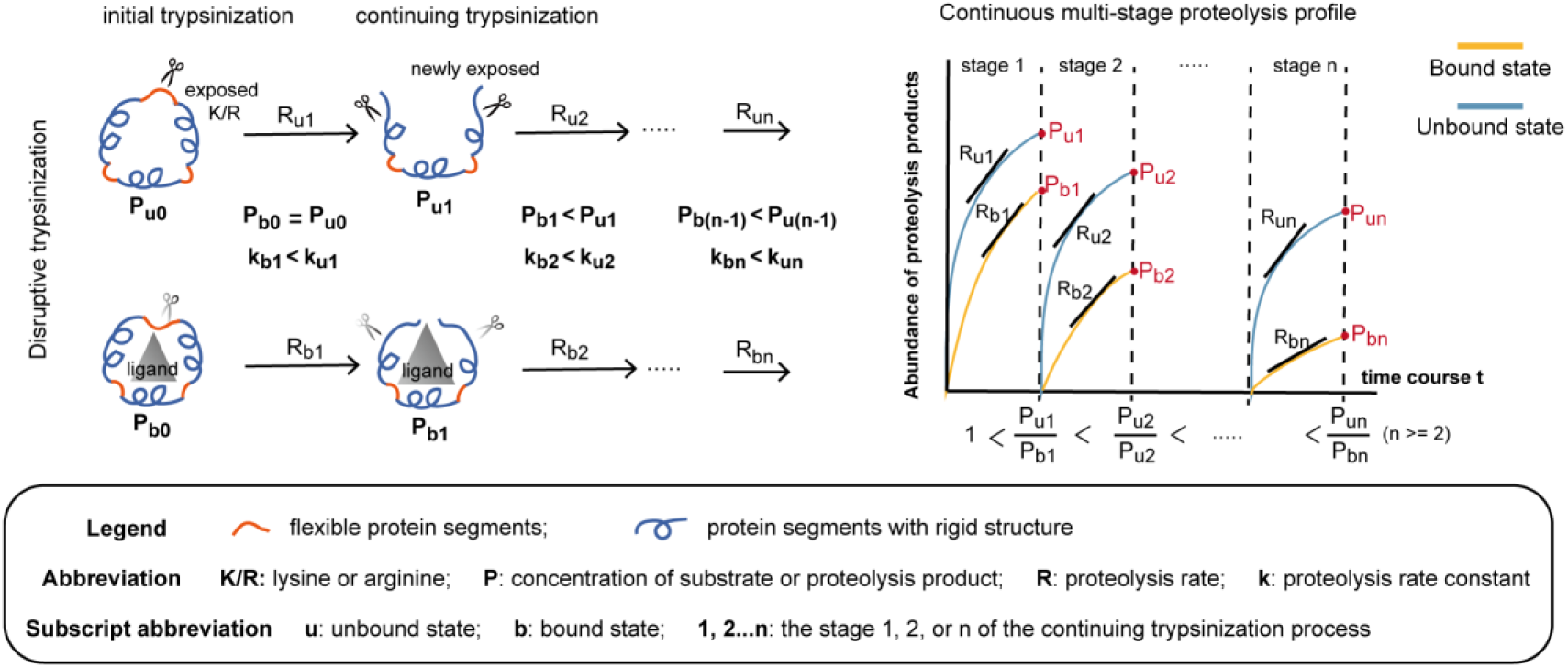
Possible mechanism for the amplified readout of protein local stability shifts in disruptive trypsinization. The overall proteolysis process for LiP-MS can be clearly separated into two steps: an initial proteolysis by proteinase K (PK) and a later denaturation-assisted complete digestion by trypsin. In contrast, the disruptive trypsinization should be conceptualized as a continuous, multi-stage proteolysis process, which comprises a successive sequential ‘steps’ (*i.e*., cleavage reactions) as each new trypsin-sensitive site is exposed. The initial proteolysis of both LiP-MS and disruptive trypsinization typically occurs at the flexible segments for their accessibility to the proteolytic sites of protease, and ligand binding will stabilize the flexible segments, thereby delaying the initial proteolysis (k_b1_ < k_u1_). In the LiP-MS procedure, the initial proteolysis is performed with a broad-specificity protease, PK, to generate large protein segments. These large protein segments in both the ligand-treated and control samples are then subjected to indiscriminate denaturation, followed by complete indiscriminate trypsinization. As such, only the proteolysis rate in the initial proteolysis is altered upon ligand binding. In disruptive trypsinization, trypsin is used for the whole multi-stage proteolysis to generate small peptides (shown in the figure). At the initial stage, only the flexible segments that contain lysine (K) or arginine (R) are cleaved due to the substrate specificity of trypsin, which results in cleaved proteins. The cleaved regions of the cleaved proteins are unstable, so they unfold and expose more K or R to facilitate continuing trypsinization. The cleaved proteins generated under the bound state are likely to retain the bound ligand for their relatively intact structures. The bound ligand could again delay the continuing trypsinization (k_b2_ < k_u2_). Moreover, due to the delayed initial trypsinization, the bound cleaved proteins also have a lower concentration than the unbound form (P_b1_ < P_u1_). According to Michaelis-Menten equation, at low substrate concentration (which is likely the case for individual proteins in the proteome sample), the digestion reaction can be regarded as a first-order reaction: proteolysis rate (R_2_) = substrate concentration (P_1_) * rate constant (k_2_). With the combination of ligand protection (k_b2_ < k_u2_) and the smaller substrate concentration (P_b1_ < P_u1_), the rate difference between bound and unbound states in stage 2 digestion is larger than that in stage 1 digestion, thus resulting in an amplified abundance difference of proteolysis products (P_u2_/P_b2_ > P_u1_/P_b1_ >1). This amplification could last until the bound ligand is dissociated from the cleaved proteins. The substrate of stage n trypsinization is the proteolysis product of stage (n-1). The slope of the proteolysis profile is defined by the proteolysis rate, R.

**Extended Data Fig. 3.**
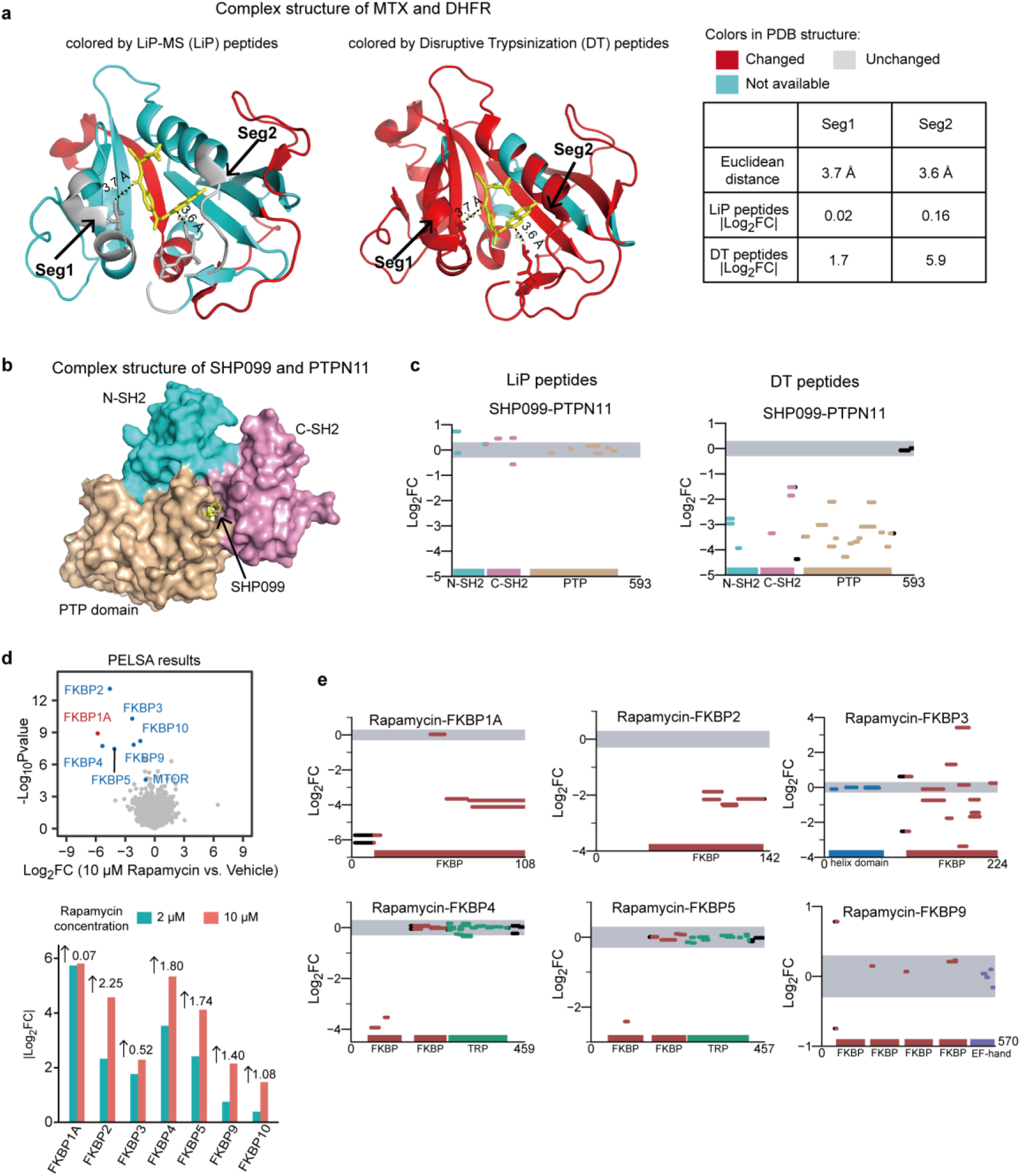
The amplified readouts in PELSA facilitate the localization of ligand-binding regions and the identification of ligand-binding proteins with low target occupancy. **a,** Complex structure of DHFR-MTX (PDB: 1U72) with protein segments colored based on the quantification results of their corresponding peptides generated by two-step digestion in LiP-MS (left) or by disruptive trypsinization in PELSA (middle): changed (-log_10_Pvalue > 2 and |log_2_FC| > 0.3), red; unchanged (|log_2_FC| < 0.3), grey; not available (not quantified or |log_2_FC| > 0.3 but -log_10_Pvalue < 2), cyan. The drug ligand MTX is shown as sticks (yellow). Two labeled segments at a distance of less than 4 Å from MTX displayed large fold changes in PELSA but remained unchanged in LiP-MS; Euclidean distance in the table is the minimal distance between the ligand atoms and the peptide atoms of the corresponding protein segment. **b,** Complex structure of PTPN11 and SHP099 (surface representation, PDB: 5EHR). PTPN11 is composed of N-SH2, C-SH2, and PTP domains; SHP099 is an allosteric inhibitor of PTPN11, known to bind at the central tunnel formed at the interface of the three domains^19^. **c,** Abundance changes of PTPN11 peptides generated by LiP-MS (left) or disruptive trypsinization (right) under 10 µM SHP099 treatment. The x axis represents the protein sequence from N to C-terminus, with protein length annotated; the y axis shows the log_2_ fold changes in abundance of the peptides (log_2_FC). The upper and lower boundaries of the grey shaded area represent log_2_FCs of 0.3 and -0.3, respectively. In LiP-MS, a number of peptides remained unchanged even though they are located in the domains associated with allosteric regulation of SHP099. By contrast, all 21 disruptive trypsinization peptides that are positioned within the domains associated with allosteric regulation of SHP099, displayed a statistically significant fold change (-log_10_Pvalue > 2 and |log_2_FC| > 0.3), whereas the 2 unchanged peptides are from the C-terminal tail of PTPN11, which do not participate in SHP099 binding. **d,** Top: volcano plot visualization of all proteins from a PELSA analysis of HeLa cell lysates exposed to 10 µM rapamycin; Bottom: comparing magnitude of fold changes (log2 transformed) of seven FKBP family proteins under 2 µM rapamycin and 10 µM rapamycin treatment. When increasing concentration of rapamycin to 10 µM, the fold change of FKBP1A remained relatively constant, while five of the remaining six FKBP family proteins showed a more than 2-fold increase in the magnitudes of fold changes (the sixth showed 1.44-fold increase; the labeled values represent log2 transformed increased values). This result indicates the low target occupancy of the remaining six FKBP family proteins under 2 µM rapamycin treatment. **e,** Local stability profiles of FKBP family proteins under 2 µM rapamycin treatment. Only peptides from FKBP domains display altered abundance.

**Extended Data Fig. 4.**
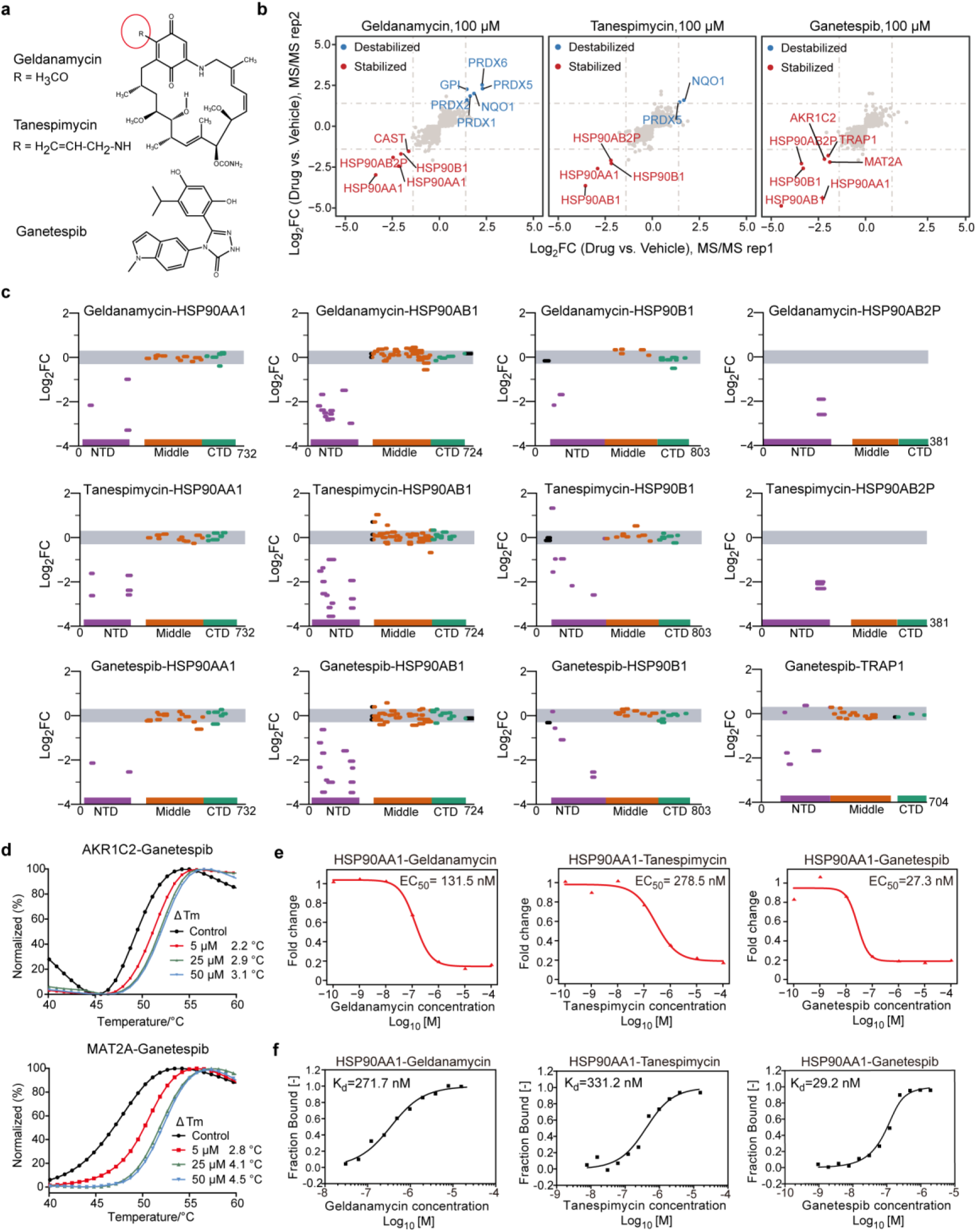
PELSA couples with dimethyl labeling quantification for reliable target protein identification, precise binding region localization, and accurate binding affinity determination. **a,** Structures of three HSP90 inhibitors used in this study. The red cycle indicates the structural difference between geldanamycin and tanespimycin. **b,** Scatter plots of protein log_2_ fold changes (**Supplementary Discussion**) in HeLa cell lysates treated with three HSP90 inhibitors in two MS/MS analyses. Proteins with |log_2_FC| >1.4 are colored as indicated in the legend. **c,** Local stability profiles of HSP90 family proteins under 100 µM geldanamycin (top), 100 µM tanespimycin (middle), and 100 µM ganetespib (bottom) treatment. NTD refers to N-terminal ATP binding domain; CTD refers to C-terminal domain. **d,** Protein melting curves of purified recombinant AKR1C2 (top) and MAT2A (bottom) after incubation with different concentrations of ganetespib. **e,** PELSA-determined dose-response curves for HSP90AA1 in HeLa cell lysates incubated with three HSP90 inhibitors at different concentrations. **f,** Dose-response curves as in (**e**) but measured by microscale thermophoresis assays using purified HSP90AA1.

**Extended Data Fig. 5.**
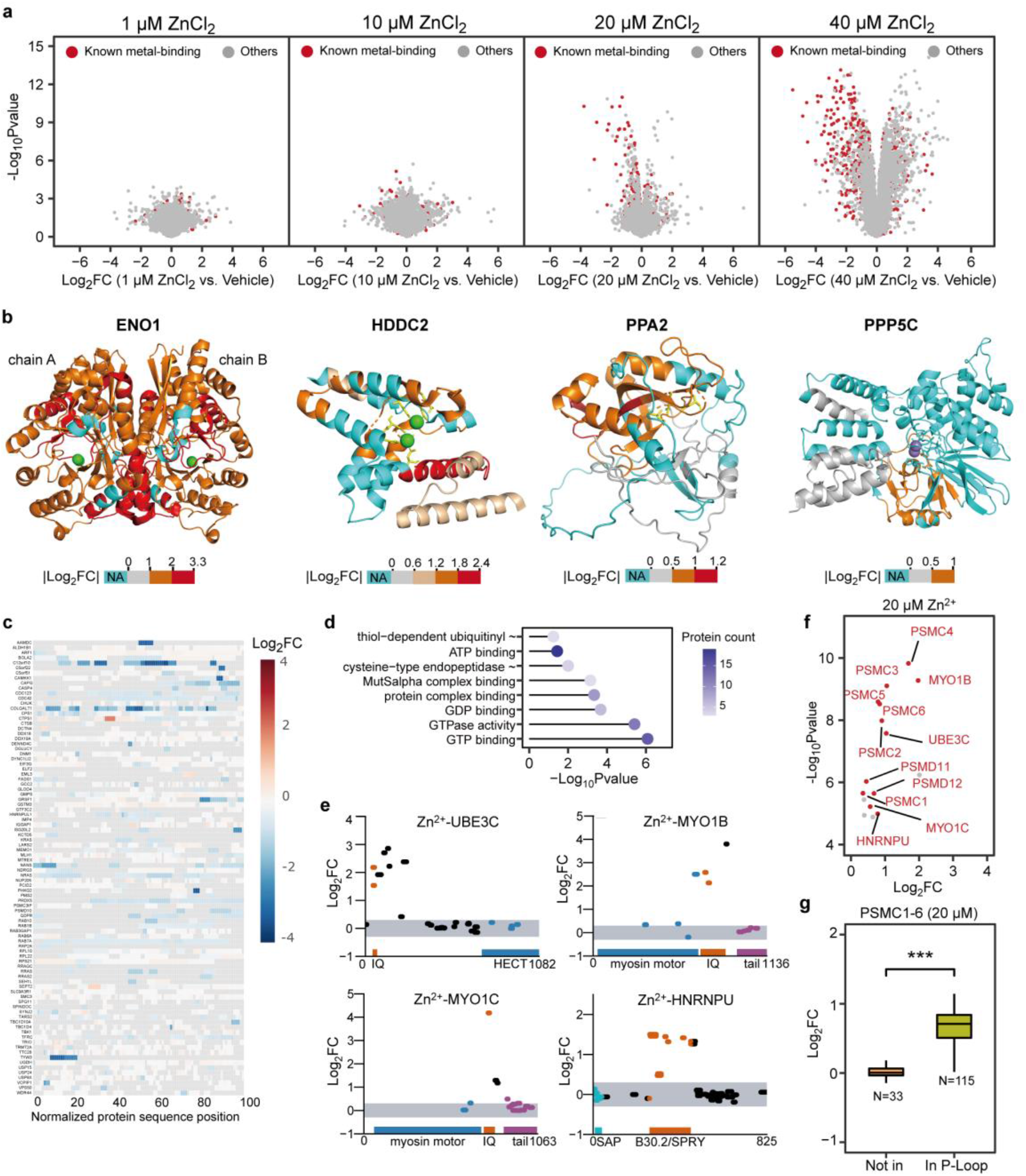
Characterization of the zinc proteome. **a,** Volcano plot visualizations of all proteins from PELSA analyses of HeLa lysates exposed to 1 μM, 10 μM, 20 μM, or 40 μM ZnCl_2_ (four lysate replicates). Uniprot-annotated known metal-binding proteins are highlighted in red. The Zn^2+^ concentrations used for further analysis are 20 μM and 30 μM (**Supplementary Discussion**). **b,** Representative Mg^2+^-binding proteins that were stabilized by 30 μM Zn^2+^. From left to right: ENO1 homodimer in complex with Mg^2+^ (PDB: 2PSN), HDDC2 in complex with Mg^2+^ (PDB: 4DMB), PPA2 (AlphaFold: AF-Q9H2U2-F1-mod), and PPP5C in complex of Mn^2+^ (PDB: 1WAO). Protein segments are colored based on log_2_FC values as indicated in the legend. NA denotes no detection (cyan). The magnesium and manganese ions are represented as green and dark-blue spheres, respectively. The Mg^2+^-binding residues are shown as yellow sticks. Mg^2+^ acts as a dimer stabilizer for enolases^21^. Therefore, global stabilization was discovered for ENO1, ENO2, and ENO3 (see also **Supplementary Table 6**). For HDHD2, PPA2 and PP5C, the Zn^2+^-induced stabilization mainly occurs at Mg^2+^-binding regions. **c,** Overview of the local stability profiles of the 90 Zn^2+^ -stabilized proteins that were not categorized as metal binding. Each row represents an individual protein with its gene name labeled on the left. Protein sequence lengths are normalized to 100. Protein segments are colored in a heatmap color mode based on their log_2_FC values (not quantified by PELSA, grey). **d,** Dot plot showing the protein counts and p values (Fisher’s exact test) of the molecular functions enriched among the 90 Zn^2+^-stabilized proteins that are not categorized as metal binding. **e,** Three IQ-motif-containing proteins (UBE3C, MYO1C, and MYO1B) were destabilized at or around IQ motifs by 30 µM Zn^2+^ treatment. IQ motifs are interacting surfaces of EF motifs^67^ which are observed stabilized by Zn^2+^ treatment. A B30.2/SPRY-domain-containing protein HNRNPU was destabilized at B30.2/SPRY-domain, which functions as a protein-interacting module in many proteins^68^. **f and g,** PELSA analysis of HeLa cell lysates treated with 20 µM Zn^2+^ (four lysate replicates). **f,** Volcano plot visualization of the top 16 most significantly destabilized proteins (log_2_FC > 0, ranked by -log_10_Pvaue, Bayes t-test). The 12 proteins destabilized at known protein-protein interaction interfaces are colored and labeled in red. **g,** Boxplots displaying the distributions of log_2_ fold changes of PSMC1-6 peptides (grouped by within and outside the P-Loop-NTPase motifs). ***p < 0.001, Wilcox signed rank test.

**Extended Data Fig. 6.**
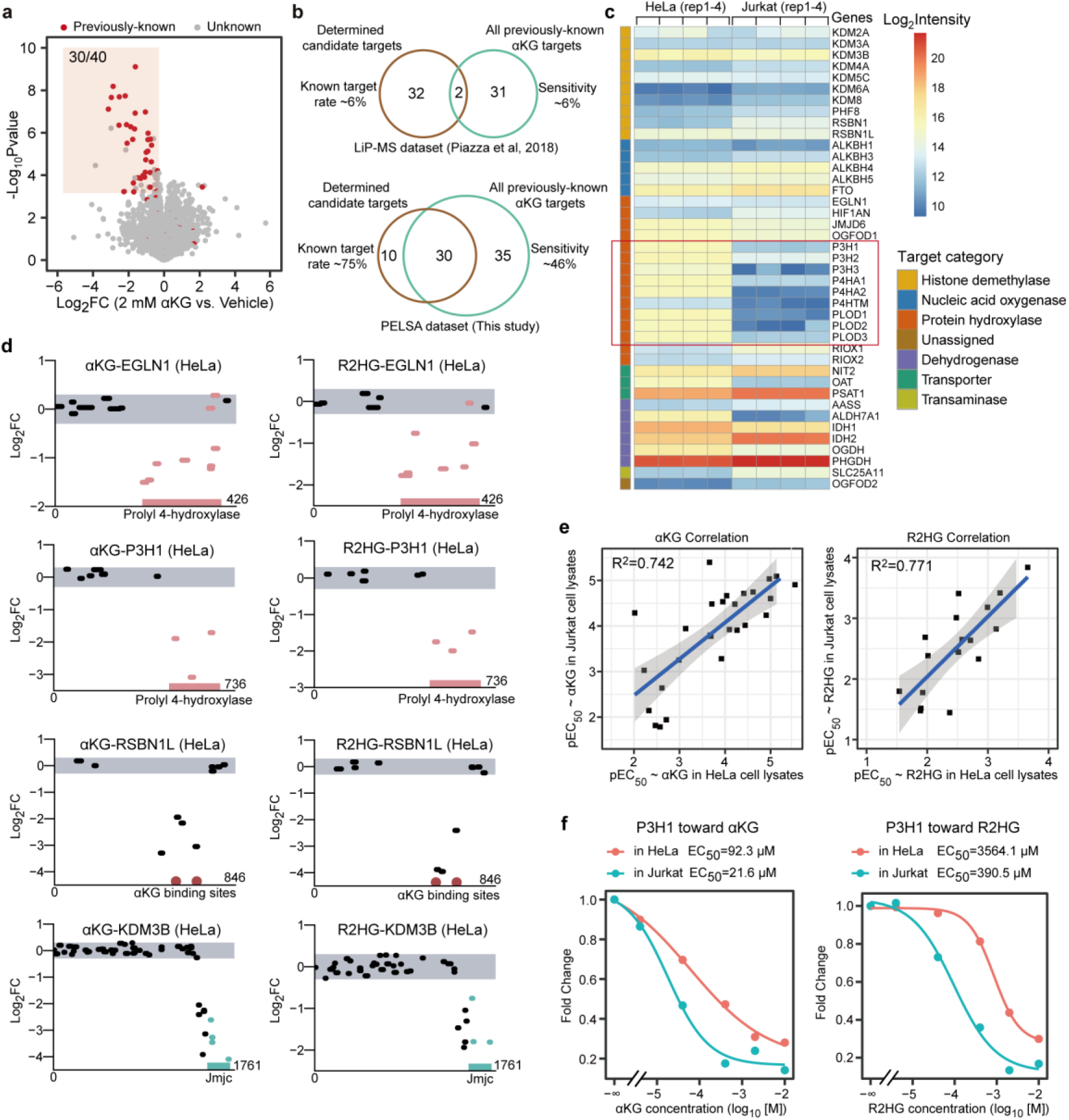
Characterization of the previously-known αKG and R2HG target proteins identified by PELSA. **a,** Volcano plot visualization of all proteins from a PELSA analysis of HeLa cell lysates exposed to 2 mM αKG (four lysate replicates). The previously-known αKG target proteins are marked in red. The left boundary and lower boundary of the red shadow denote log_2_FC of -0.5 and - log_10_Pvalue of 3.4, respectively. Among the 40 candidate target proteins, 30 are previously-known αKG target proteins. **b,** Venn plots showing the numbers of candidate αKG target proteins determined by LiP-MS or PELSA (brown cycle) and all previously-known αKG target proteins included in the LiP-MS or PELSA dataset (green cycle)^52^. Known target rate denotes the percentage of previously-known αKG target proteins among all determined candidate αKG target proteins; sensitivity denotes the percentage of previously-known αKG target proteins that were determined as candidate αKG target proteins (true positive / true positive + false negative). **c,** Comparing protein abundance of the previously-known αKG target proteins between HeLa cells and Jurkat cells (four lysate replicates). **d,** Local stability profiles of representative previously-known αKG targets in HeLa cell lysates by 10 mM αKG treatment (left) and 10 mM R2HG treatment (right). **e,** Pearson correlation between pEC_50_ values determined in PELSA analyses of HeLa and Jurkat cell lysates. Top, αKG toward previously-known αKG target proteins; Bottom, R2HG toward previously-known αKG target proteins. **f,** Dose-response curves of P3H1 measured by PELSA in HeLa and Jurkat cell lysates toward αKG and R2HG.

**Extended Data Fig. 7.**
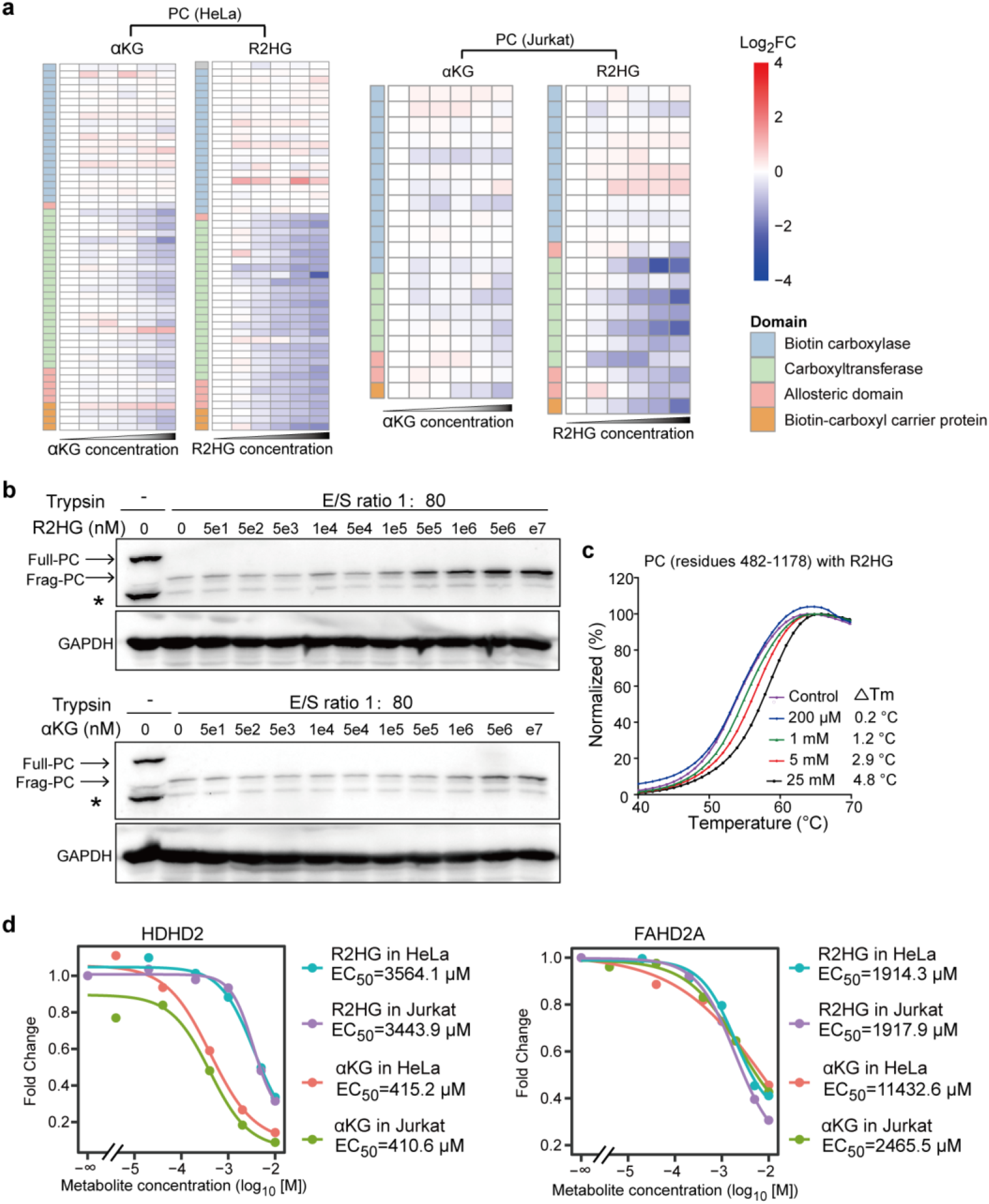
Characterization of the previously-unknown αKG and R2HG target proteins. **a,** Comparing local affinity profiles of PC toward αKG (left) and R2HG (right) in the HeLa and Jurkat cell lysates. **b,** Western-blot readout confirms the higher affinity of PC toward R2HG than αKG. **c,** Protein melting curves for the purified recombinant PC segment (residues 482-1178) under different concentrations of R2HG. **d,** Binding affinities of HDHD2 (left) and FAHD2A (right) toward αKG and R2HG in HeLa and Jurkat cell lysates. HDHD2 and FAHD2A are two proteins without known functions.

**Extended Data Fig. 8.**
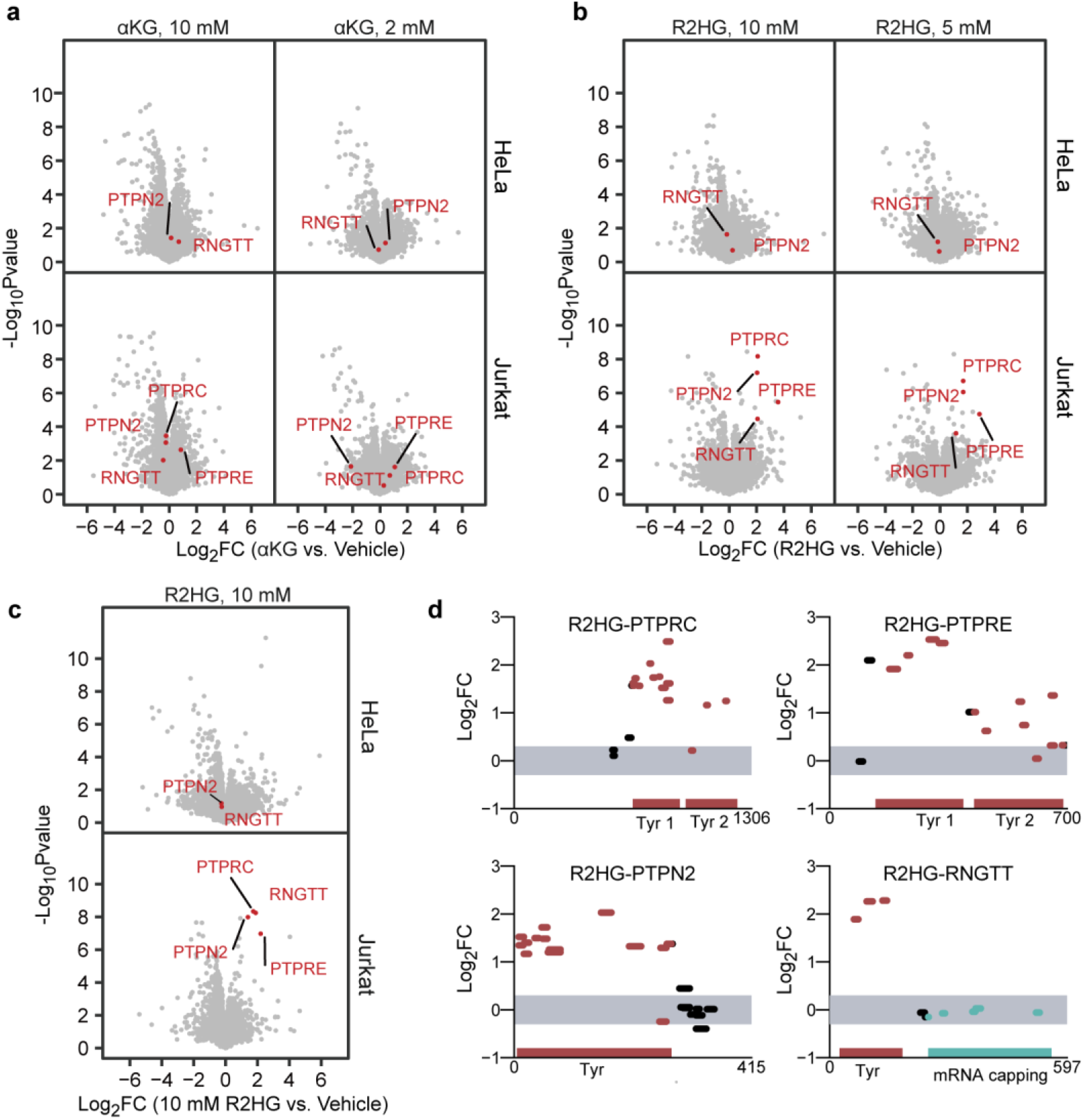
Characterization of four tyrosine-phosphatase-domain-containing proteins identified by PELSA. **a,** Volcano plot visualizations of all proteins from PELSA analyses of HeLa (top) and Jurkat (bottom) cell lysates exposed to 10 mM (left) and 2 mM (right) αKG (four lysate replicates). Four tyrosine-protein phosphatase domain-containing proteins (PTPRC, PTPN2, PTPRE, and RNGTT) stay unchanged by either 10 mM or 2 mM αKG treatment. **b,** Volcano plot visualizations as in (**a**), but for 10 mM (left) and 5 mM (right) R2HG treatment. PTPRC, PTPN2, PTPRE, and RNGTT are only destabilized by R2HG in Jurkat cell lysates. **c,** A biological replicate of PELSA analysis on HeLa and Jurkat cell lysates treated with 10 mM R2HG confirmed the R2HG-induced Jurkat-specific destabilizations of PTPRC, PTPN2, PTPRE, and RNGTT. Results represent four lysate replicates per PELSA analysis. **d,** A biological replicate of PELSA R2HG analysis in Jurkat cell lysates confirmed that R2HG-induced destabilizations of PTPRC, PTPN2, PTPRE, and RNGTT occur at their Tyr domains. Tyr is an abbreviation of protein tyrosine phosphatase domain.

